# A MALDI-TOF assay identifies nilotinib as an inhibitor of inflammation in acute myeloid leukaemia

**DOI:** 10.1101/2021.03.29.437557

**Authors:** José Luis Marín-Rubio, Rachel E. Heap, Tiaan Heunis, Maria Emilia Dueñas, Joseph Inns, Jonathan Scott, A. John Simpson, Helen Blair, Olaf Heidenreich, James M. Allan, Barbara Saxty, Matthias Trost

## Abstract

Inflammatory responses are important in cancer, particularly in the context of monocyte-rich aggressive myeloid neoplasm. We developed a label-free cellular phenotypic drug discovery assay to identify anti-inflammatory drugs in human monocytes derived from acute myeloid leukaemia (AML), by tracking several biological features ionizing from only 2,500 cells using matrix-assisted laser desorption/ionization-time of flight (MALDI-TOF) mass spectrometry. A proof-of-concept screen showed that the BCR-ABL inhibitor nilotinib, but not the structurally similar imatinib, blocks inflammatory responses. In order to identify the cellular (off-)targets of nilotinib, we performed thermal proteome profiling (TPP). Unlike imatinib, nilotinib and other later generation BCR-ABL inhibitors inhibit the p38α-MK2/3 signalling axis which suppressed the expression of inflammatory cytokines, cell adhesion and innate immunity markers in activated human monocytes derived from AML. Thus, our study provides a tool for the discovery of new anti-inflammatory drugs, which could contribute to the treatment of inflammation in myeloid neoplasms and other diseases.

**Key Points:** Label-free cell-based assay identifies new anti-inflammatory drugs using MALDI-TOF MS. Nilotinib reduces inflammation by inhibition of MAPK14-MK2/3 signalling axis in AML.

## INTRODUCTION

Monocyte-derived macrophages play important roles in both physiological and pathological processes. The activation of these cells during pathogen infection or other insults leads, in part, to the pathophysiology of inflammation (Gordon & Taylor, 2005). Monocytes can recognize various damage-associated molecular (DAMPs) and pathogen-associated molecular patterns (PAMPs) through specific receptors, such as Toll-like receptors (TLRs). Activation of these receptors leads to the production of cytokines, chemokines, and mediators that are involved in inflammation in myeloid leukemia (Binder *et al*, 2018; Craver *et al*, 2018; Stuhlmuller *et al*, 2000).Several hematopoietic disorders, including lymphoproliferative disorders and myelodysplastic syndromes, which possess a high risk of transformation to leukaemia, have been linked to aberrant TLR signalling (Craver *et al*., 2018; Hemmati *et al*, 2017; Monlish *et al*, 2016). Moreover, TLR2 and TLR4 show significantly higher expression in the bone marrow of patients with myeloid leukemia (Monlish *et al*., 2016). TLR activation leads to intracellular signalling cascades including the mitogen-activated kinase (MAP) pathways (Kim & Choi, 2010; Suzuki *et al*, 2018), increasing the expression and secretion of inflammatory cytokines and chemokines including interleukins, interferons, and tumour necrosis factor alpha (TNF-α) (Mogensen, 2009). A shared characteristic of many hematologic malignancies is the overproduction of inflammatory cytokines, particularly TNF-α and IL-6 in myeloid malignancies (Craver *et al*., 2018).

The discovery of drugs that prevent chronic or acute inflammation is a major goal of the pharmaceutical industry for the treatment of infectious diseases, autoimmune diseases, and cancer (Inglese *et al*, 2007). The discovery and design of new compounds for inhibiting inflammation are typically achieved through high-throughput screening (HTS) approaches. As such, there is a growing need to develop physiological phenotypic HTS assays that can be used for monocytic cell lines and primary cells to accelerate the discovery of inhibitors (Dulai & Sandborn, 2016; Li *et al*, 2017). Mass spectrometry (MS)-based readouts in drug discovery have been largely dominated by instruments comprising of solid-phase extraction (SPE) coupled electrospray ionization (ESI; i.e. RapidFire) or surface-based MS techniques, such as matrix-assisted laser/desorption ionization (MALDI) (McLaren *et al*, 2021). MALDI-time of flight (MALDI-TOF) MS is a versatile, label-free technique that has the potential to accelerate HTS of promising drug candidates (De Cesare *et al*, 2018; Guitot *et al*, 2017; Heap *et al*, 2017). MALDI-TOF MS is tolerant to a number of standard buffer components and has rapidly become popular in the field of HTS drug discovery due to its versatility, requiring very small sample quantities, and minimal sample clean-up (Chandler *et al*, 2017; Dreisewerd, 2014; Guitot *et al*., 2017; Heap *et al*., 2017; Ritorto *et al*, 2014; Wang *et al*, 2008). MS-based screening approaches offer the possibility to simultaneously track several molecules in a label-free manner, provide excellent signal-to-noise, reproducibility, assay precision, and a significantly reduced reagent cost when compared to fluorescence-based assays. Furthermore, fluorescence and chemiluminescence methodologies in primary cells and other read-outs, such as antibody-based assays, are very expensive, making full-deck screens of millions of compounds difficult (Adegbola *et al*, 2018; McLaren *et al*., 2021; Neefjes & Dantuma, 2004; Wang *et al*., 2008). MALDI-TOF MS, similar to acoustic mist ionization mass spectrometry (Smith *et al*, 2021), requires little sample preparation, allows rapid screening and it has high specificity and sensitivity (McLaren *et al*., 2021). Whole cell analyses or cellular assays for evaluating compound efficacy affecting a cellular phenotype present an interesting challenge for MALDI-TOF MS analysis as the system becomes inherently more complex. One of the attractive qualities of this type of assay for the pharmaceutical industry is that the cellular assays provide biologically relevant information about the physiology and general condition of a cell, such as cell viability and protein activity in a single cellular screen (Butcher, 2005; Michelini *et al*, 2010). This simultaneous acquisition of information is known as multiplexing and is desirable in cellular assays, as it significantly increases the screening efficiency of the compound library (Gerets *et al*, 2011).

In this study, we have developed a cellular MALDI-TOF MS assay that is highly reproducible, robust, and sensitive in different cell lines, with which we can identify two main signatures altered under different pro-inflammatory stimuli. Using this MALDI-TOF MS assay, we performed a blind screen of 96 compounds to assess their potential anti-inflammatory effects on human monocytes derived from acute myeloid leukaemia (AML). We discovered that nilotinib, but not the structurally related imatinib, inhibits the pro-inflammatory phenotype. Using thermal proteome profiling (TPP) (Becher *et al*, 2016; Franken *et al*, 2015; Jarzab *et al*, 2020; Mateus *et al*, 2020; Mateus *et al*, 2016; Miettinen *et al*, 2018; Reinhard *et al*, 2015; Savitski *et al*, 2014; Zinn *et al*, 2021), and whole proteome analyses we identified direct targets and downstream regulation of nilotinib during inflammation events. Our data showed that nilotinib and other 2^nd^ and 3^rd^ generation BCR-Abl inhibitors, but not imatinib, are capable to block the p38α MAP kinase signalling pathway, which abates monocyte activation and differentiation, reduces cytokine release, and prevents inflammation in AML.

## RESULTS

### A novel MALDI-TOF MS cellular assay to characterize inflammatory phenotypes

Recently, high-throughput MALDI-TOF MS has been used successfully for *in vitro* assays of specific enzymes (De Cesare *et al*., 2018; De Cesare *et al*, 2020; Guitot *et al*., 2017; Heap *et al*., 2017). Here, we tested if MALDI-TOF MS can be used for phenotypic assays to monitor inflammatory responses by identifying differences in the “fingerprint” of biomolecules ionized when whole cells are spotted onto the MALDI target (**Figure 1A**). As a proof of concept, we tested if we could distinguish phenotypes of THP-1 cells, a model of human monocytes derived from AML (Bosshart & Heinzelmann, 2016), upon stimulation with a number of pro-inflammatory stimuli, namely LPS, Pam_2_CSK_4_, Pam_3_CSK_4_, polyinosinic:polycytidylic acid (poly(I:C)), and polyadenylic-polyuridylic acid (poly(A:U)), which activate TLR4, TLR2/6, TLR2/1, and TLR3, respectively, as well as interferon-γ (IFN-γ). Principal component analysis (PCA) of all biomolecules detected in the range of *m/z* 2,000-20,000 showed clear separation of monocytes treated with TLR-agonists, while IFN-γ treatment did not separate from untreated, suggesting that the features are not dependent on interferon-stimulation but rather TLR dependent (**Figure 1B**). A loading plot analysis of the PCA results identified the features at *m/z* 4632,4964, and 6891 as the main drivers for the difference in the phenotypes (**Figure 1C**). The features at *m/z* 4632 and 6891 were significantly reduced, while *m/z* 4964 increased in LPS-stimulated cells (**Figure 1D-E** and **Supplementary Figure 1A**). As the feature at *m/z* 6891 showed a greater variability between replicates, we selected the features m/z 4632 and 4964 for further analysis of the phenotypes. We confirmed that other TLR ligands such as Pam_2_CSK_4_ and Pam_3_CSK_4_, which activate TLR1/2 and TLR2/6, respectively, did also increase this ratio and in a dose-dependent manner, while interferon-gamma (IFN-γ), poly(I:C) and poly(A:U) did not affect it (**Figure 1F-G** and **Supplementary Figure 1A**). Thus, we used the feature at *m/z* 4632 as a resting phenotype biomarker and feature *m/z* 4964 as an LPS-stimulated monocyte phenotype biomarker. To verify that this phenotype was not due to cell death, we treated the cells with 0.5 µM staurosporine for 24h before analysis with our MALDI-TOF assay (**Figure 2D**) and cell viability assay (**Supplementary Figure 1L**). Together, these results indicate that biomarkers identified by MALDI-TOF MS can be used to detect changes in the inflammatory phenotype downstream of TLRs in AML cells.

**Figure 1.**
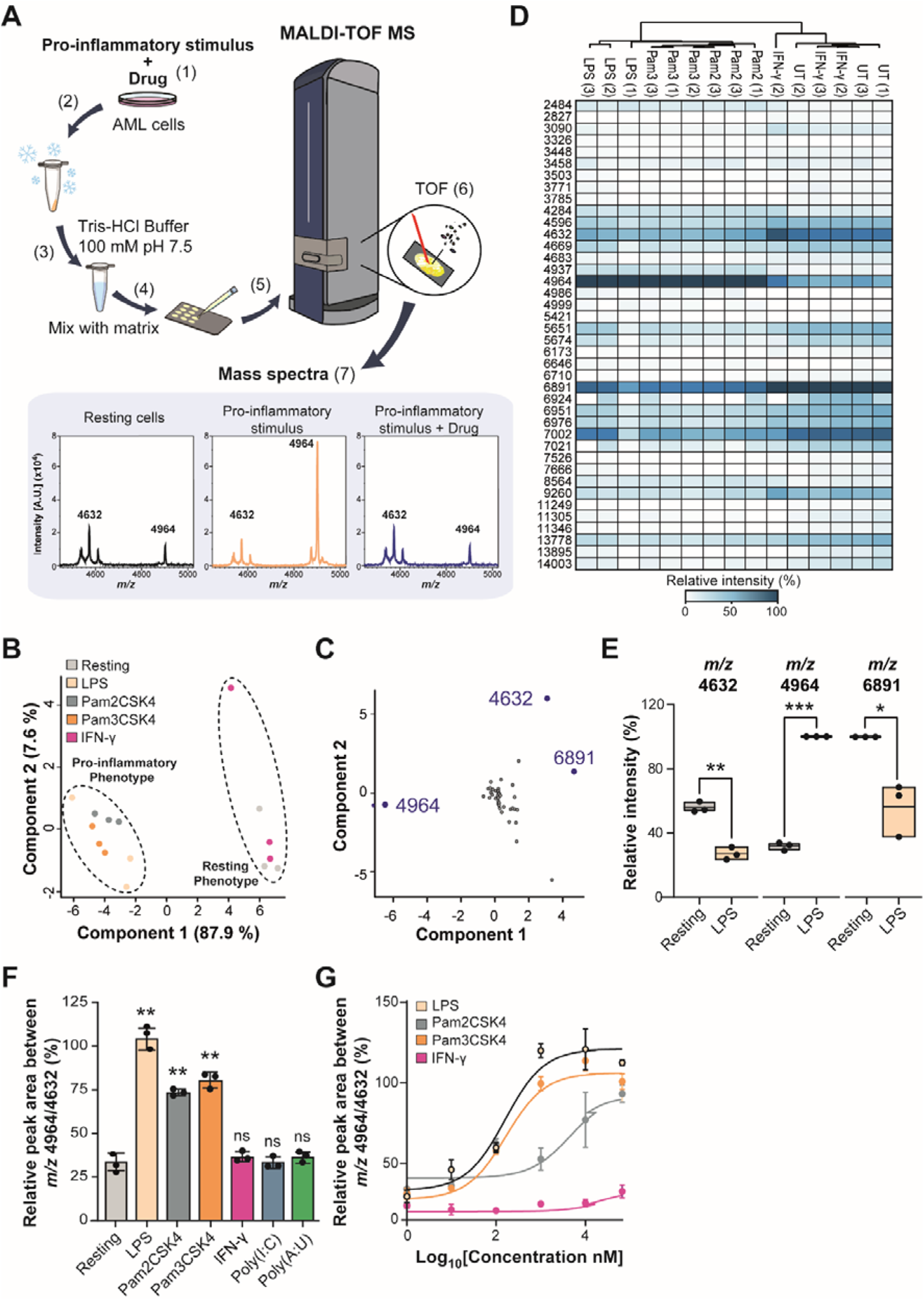
Identification of features associated with the monocyte inflammatory phenotype by MALDI-TOF MS. **A)** Workflow of the MALDI-TOF MS assay: (1) AML cells were pre-treated with a drug for 1h before adding a pro-inflammatory stimulus for 24h. (2) Cells were frozen on dry ice, thawed and (3) washed with 100 mM Tris-HCl buffer at 4°C. (4) Cells were mixed with a matrix (10 mg/mL α-cyano-4-cinnamic acid in 50% acetonitrile, 0.1% trifluoroacetic acid). (5) Cells were analyzed in a rapifleX PharmaPulse MALDI TOF mass spectrometer. (6) In the ionization chamber a laser is used to produce ions in the gas phase. These ions are separated according to their time-of-flight (TOF) in a field-free region. The smaller ions reach the detector first, followed by the bigger ions, according to the *m/z* ratio. (7) The detector converts the received ions into electrical current which is amplified and digitized in m/z spectra. **B)** Unsupervised PCA plot of LPS, Pam_2_CSK_4_, Pam_3_CSK_4_ and IFN-γ treated cells showing separation of cells treated with bacterial ligands treated and resting monocytes. **C)** Loading plot derived from PCA in B) showing that m/z 4964 and 4632 contribute predominantly to the separation of the two clusters in component 1 and component 2. **D)** Unsupervised heat map of the relative intensities of three biological replicates of THP-1 cells treated with 100 ng/mL LPS, Pam_2_CSK_4_, Pam_3_CSK_4_, 100 U/mL IFN-γ, 1 μg/mL poly(I:C) and poly(A:U) for 24h compared to resting cells **E)** Box plots of significantly changing intensities between resting and LPS-treated monocytes identified at *m/z* 4632 and 4964. **F)** Relative quantitation from three biological replicates of THP-1 cells treated with 100 ng/mL LPS, Pam_2_CSK_4_, Pam_3_CSK_4_ and 100 U/mL IFN-γ for 24h compared to resting cells. **G)** Titration of LPS, Pam_2_CSK_4_ and Pam_3_CSK_4_-treated cell from 10 - 100 ng/mL of stimulus. Significant differences between two groups were determined by Mann-Whitney U-test. The statistical significance of the comparisons with resting is indicated as follows: ns, not significant; ***, *P* ≤ 0.001; *, *P* ≤ 0.05. Error bars represent the standard deviation of three biological replicates.

**Figure 2.**
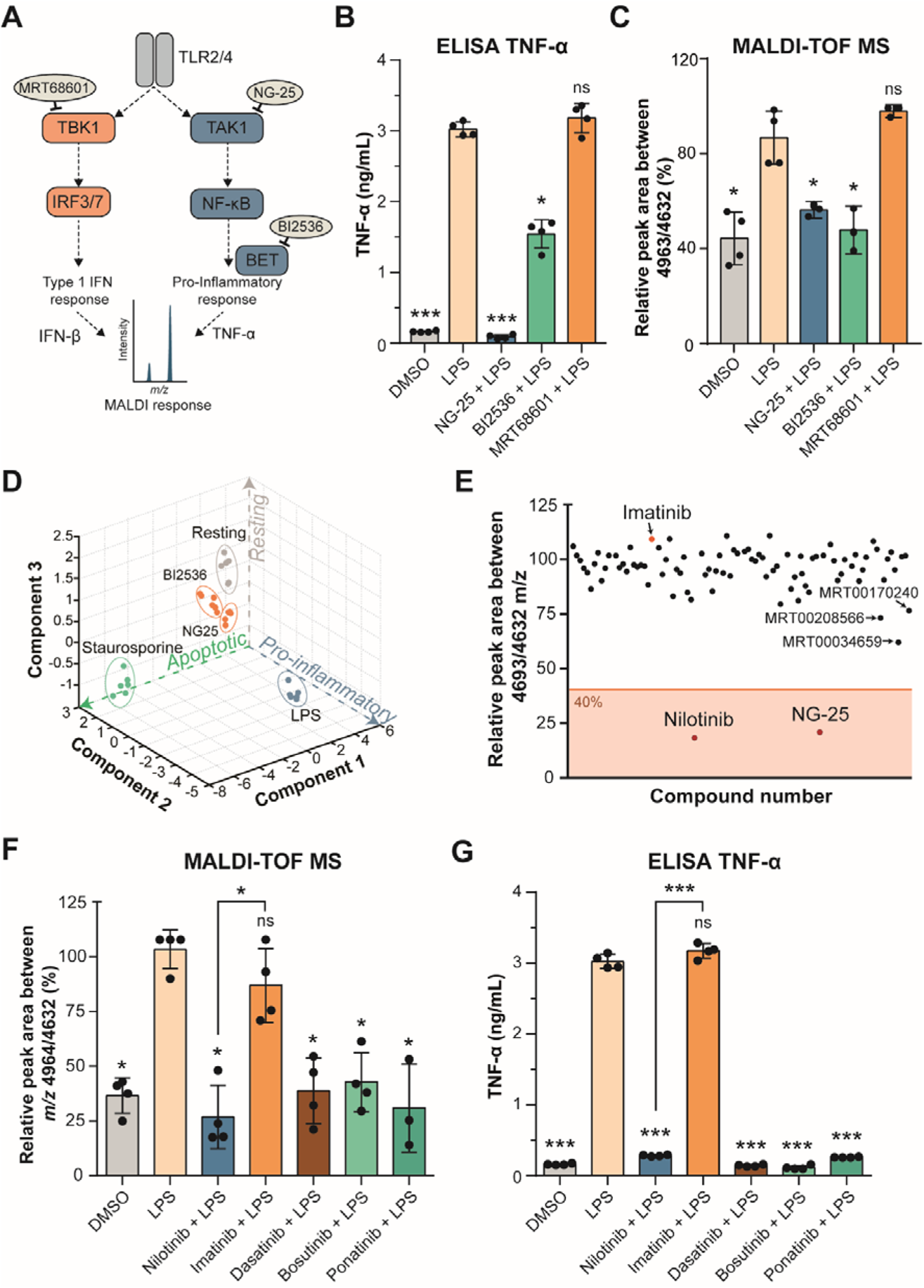
Nilotinib is a positive hit in a cellular MALDI-TOF MS drug discovery screen and inhibits inflammation in response to LPS. **A)** Inhibition of the TLR2 and TLR4 signalling pathways. MRT68601 inhibits TBK1 (TANK-Binding Kinase-1) which blocks production of type I interferons (IFNs). NG-25 inhibits TAK1 (TGF-Beta-Activated Kinase-1) which induces TNF-α and IL-6 production. BI2536 inhibits BET proteins (Bromodomain and Extra-Terminal motif proteins) which are required for pro-inflammatory gene transcription. **B)** TNF-α secretion of THP-1 cells treated in either vehicle control-treated (DMSO), 100 ng/mL LPS-treated, or pre-treated with 5 µM NG-25, BI2536 or 1 µM MRT68601 for one hour before 100 ng/mL LPS-treatment for up to 24h measured by ELISA. **C)** MALDI-TOF MS relative quantitation of the normalized ratio *m/z* 4964/4632 in THP-1 cells as treated in B). **D)** PCA of all features identified by the MALDI-TOF MS of experiment in B), and 0.5 µM staurosporine-treated cells. **E)** Compound hit map of the mean of three biological replicates with 40% effectiveness cut off showing nilotinib and NG-25 as positive hits and imatinib as negative hit (arrows). **F)** MALDI-TOF MS relative quantitation of the normalized ratio *m/z* 4964/4632 in THP-1 cells in vehicle control-treated (DMSO), 100 ng/mL LPS-treated, or pre-treated with 5 µM nilotinib, imatinib, 1 µM dasatinib, bosutinib or ponatinib for one hour before 100 ng/mL LPS-treatment for up to 24h. **G)** TNF-α secretion of cells treated as in F.) measured by ELISA after 24h. Significant differences between o groups were determined by a Mann-Whitney U-test. The statistical significance of the comparisons with LPS is indicated as follows: ns, not significant; *, *P* ≤ 0.05; **, *P* ≤ 0.01; ***, *P* ≤ 0.001. Error bars represent the standard deviation of four biological replicates.

### The MALDI-TOF MS cellular assay can detect inhibitors of inflammation

Next, we used NG-25, a TAK1 inhibitor which blocks TNF-α production; BI2536, a PLK1 inhibitor that has been show to inhibit pro-inflammatory gene transcription due to inhibition of BET proteins (Malik *et al*, 2015); as well as MRT68601, a highly selective and potent TBK1 inhibitor which blocks production of type I interferons (IFNs) (Newman *et al*, 2012; Xu *et al*, 2020), to test if pharmacological inhibition of these pathways can be detected in our assay (**Figure 2A**). As LPS provided the greatest response of all inflammatory stimuli (**Figure 1F-G**), this stimulus was used for further assays. THP-1 cells were pre-treated with the inhibitors for one hour before LPS-stimulation. NG-25 and BI2536 treatment, but not MRT68601, reduced TNF-α secretion (**Figure 2B**). It has been published that MRT68601 suppresses the production of type I interferons but increases production of pro-inflammatory cytokines (Xu *et al*., 2020), which we could not confirm within our experiments. Comparably to the cytokine secretion, our MALDI-TOF MS assay showed that treatment with NG-25 or BI2536 blocked the inflammatory phenotype and was indistinguishable to vehicle control cells not treated with LPS, while MRT68601-treated cells still showed an increased ratio at *m/z* 4964/4632, similar to LPS-treated cells (**Figure 2C** and **Supplementary Figure 1B**). To verify that this phenotype was not due to cell death, we treated the cells with 0.5 µM staurosporine for 24h before analysis with our MALDI-TOF assay. Our results showed that apoptosis leads to a distinct cluster in the PCA compared to control or LPS-treated cells (**Figure 2D**, suggesting that this assay can be multiplexed to identify other phenotypes such as cell toxicity. Moreover, using all biomolecule features, PCA analysis showed that treatment with NG-25 and BI2536 reverted the fingerprint back close to control (**Figure 2D**). This data shows that pharmacological intervention of inflammatory pathways can be detected by our assay and that NG-25 can be used as a positive control compound in a high-throughput screen.

### Label-free MALDI-TOF MS identifies nilotinib as an anti-inflammatory compound

Once the analytical workflow was established to identify an inflammatory phenotype in THP1 cells, we evaluated the potential of the assay to discriminate positive and negative inhibitors of inflammation in a proof-of-concept blind screen of 96 compounds (**Supplementary Table 1**). The assay showed very good sensitivity (z’=~0.8) and reproducibility (R^2^ > 0.8) (**Supplementary Figure 2A-B**). Only two positive hits were observed when applying a cut-off of 40% to the ratio *m/z* 4964/4632, as previously determined for MALDI-TOF assays (De Cesare *et al*., 2020; Heap *et al*., 2017). Unblinding revealed that these compounds were NG-25 and nilotinib (Tasigna^®^) (**Figure 2E**). Nilotinib is a BCR-ABL tyrosine kinase inhibitor (Pan *et al*, 2019; Wei *et al*, 2010) and is a second-generation derivative of imatinib (Gleevec^®^) (Pan *et al*., 2019), which was also included in the panel screened. However, imatinib did not affect the phenotype of inflammatory monocytes, while nilotinib-treated monocytes did not display an LPS-induced inflammatory phenotype and showed no differences compared to non-activated monocytes in the ratio of *m/z* 4964/4632 (**Figure 2F** and **Supplementary Figure 1B**). Moreover, we tested by MALDI-TOF MS other tyrosine kinase inhibitors (TKIs) used in CML patients (Pophali & Patnaik, 2016): the second-generation TKI, dasatinib and the third-generation TKIs, bosutinib and ponatinib, which all also reduced the *m/z* 4964/4632 ratio and thus, inflammatory responses (**Figure 2F** and **Supplementary Figure 1C**) without affecting cell viability (**Supplementary Figure 1M**)

In order to validate the MALDI-TOF MS results, we measured secreted TNF-α levels in THP-1 cells treated with each inhibitor. Nilotinib, as well as treatments with the second and third generation TKIs reduced expression and secretion levels of TNF-α, while imatinib did not affect TNF-α levels upon LPS treatment (**Figure 2G**). Both MALDI-TOF MS and TNF-α ELISA data provided a similar IC50 of approximately 400-500 nM for the inhibitory effect of nilotinib (**Supplementary Figure 2C-D**). This data indicates that nilotinib and later generation TKIs are able to block inflammatory responses downstream of TLR activation, while the structurally similar imatinib cannot.

### Nilotinib blocks the inflammatory response by inhibiting the p38α MAP kinase pathway

Nilotinib is known to inhibit BCR-ABL and other tyrosine kinases.(Pan *et al*., 2019) A putative role of these kinases in inflammatory signalling downstream of TLRs is unknown. Moreover, the target involved in this particular function is likely a kinase and must be a specific target for nilotinib, since the treatment with imatinib did not result in the same phenotype. In order to identify the off-targets responsible for the nilotinib-specific anti-inflammatory phenotype, we performed thermal proteome profiling (TPP) by multiplexed quantitative mass spectrometry using tandem mass tags (TMT) (**Figure 3A**, **Supplementary Table 2**, and **Supplementary Table 3**). This method is based on determining the changes in the thermal stability of proteins, which may be due to direct drug binding, drug-induced conformation changes, binding to other cellular components, or post-translational modifications such as phosphorylation (Becher *et al*., 2016; Franken *et al*., 2015; Mateus *et al*., 2020; Miettinen *et al*., 2018; Saei *et al*, 2021; Savitski *et al*., 2014). We pre-treated THP1 cells for 1 h with nilotinib or imatinib before stimulation with LPS for 15 min. We reduced the LPS treatment to 15 min in order to avoid changes in protein abundance due to transcriptional responses to LPS activation. Overall, we identified 5,565 proteins in our TPP analysis (**Supplementary Table 4** and **Supplementary Figure 3**). Only seven proteins changed significantly in all four replicates in melting temperature between nilotinib and imatinib with a standard deviation below two degrees. For three of these proteins, we detected increased ΔT_m_ in nilotinib-treated samples. The strongest hit was MAP kinase p38α (MAPK14) with an increased ΔT_m_ of 6°C in nilotinib-treated samples (**Figure 3B**). The closely related isoform p38γ (MAPK12) did not show significant differences (**Supplementary Figure 4A**). Also, MAPK activated protein kinase 3 (MK3), a downstream protein target of p38, showed an increased ΔT_m_ of 2.5°C in nilotinib-treated samples (**Figure 3C**). The remainder of significant proteins are shown in **Supplementary Figure 4B-F**. To validate p38 alpha, we performed western blots of lysates from a cellular thermal shift assay. The thermal denaturation temperatures for p38 alpha in DMSO and LPS-treated monocytes were 50.3°C and 50.2°C, respectively. In accordance with the TPP data, the thermal denaturation temperatures for p38 with nilotinib was 60.7°C, while with imatinib it was 50.8°C (**Figure 3D-E**). Examination of the TPP data for the upstream mitogen-activated protein kinase kinase 3 (MKK3), MKK4 and MKK6 (**Supplementary Figure 4G-J**) did not show any differences. We further performed a concentration compound range experiment at 56°C to determine the amounts of nilotinib required to alter protein thermal stability of p38. We found that nilotinib was able to stabilize p38 at an EC50 concentration of 4.6 µM (**Supplementary Figure 4K-L**). Moreover, we found that nilotinib is also able to affect p38 stability without LPS (**Supplementary Figure 4M-N**).

**Figure 3.**
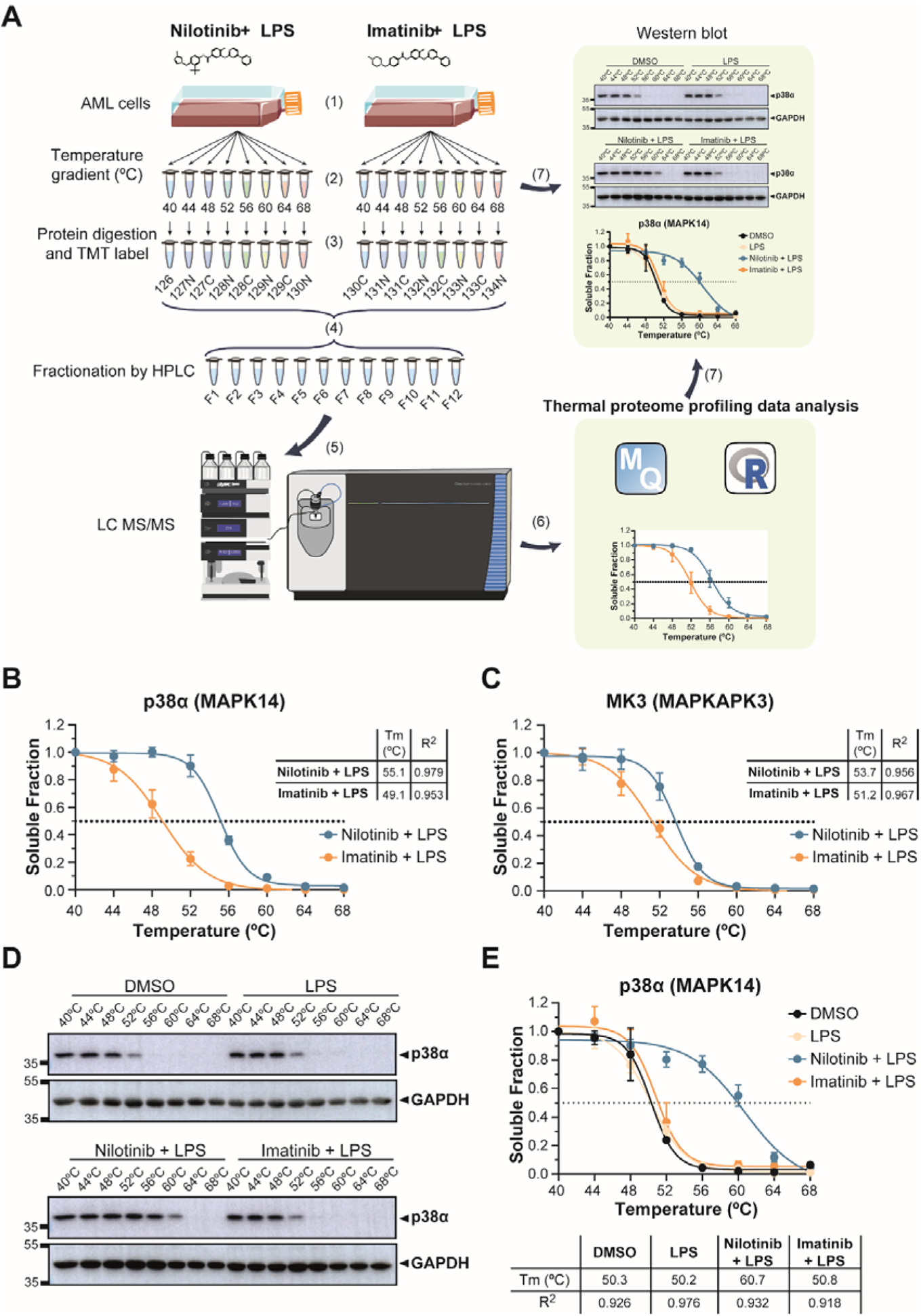
Thermal proteome profiling (TPP) reveals stabilization of p38 alpha and MK3 in nilotinib-treated compared to imatinib-treated cells. **A)** Workflow of thermal proteome profiling (TPP): (1) AML cells were pre-treated with drug for 1h before adding LPS for 15 min. (2) Cells were split in 8 different tubes and heated for 3 min at a range of temperatures (40, 44, 48, 52, 56, 60, 64 and 68°C). Cells were lysed by freeze-thawing and precipitates removed by ultracentrifugation. (3) Proteins in the supernatants were digested with trypsin, labelled with 16-plex tandem mass tags (TMT) and (4) mixed. TMT-labelled peptides were fractionated using high-performance liquid chromatography (HPLC). (5) Peptides were analysed in an Orbitrap Fusion Lumos Tribrid mass spectrometer. (6) TPP data analysis was performed using MaxQuant and the computing environment R. (7) To validate the results, target proteins were analysed by western blot and melting temperatures (Tm) were calculated. **B)** Determination of the thermostability of p38 alpha (MAPK14), and **C)** MK3 (MAPKAPK3) at the indicated temperatures with 5 µM nilotinib or imatinib for one hour before 100 ng/mL LPS-treatment for up to 15 min by TPP analysis. Table insert shows melting temperature (Tm °C) and R^2^. **D)** Determination of the thermostability of p38 alpha at the indicated temperatures in vehicle control (DMSO), 100 ng/mL LPS and pre-treated with 5 µM nilotinib or imatinib for one hour before 100 ng/mL LPS-treatment for up to 15 min. GAPDH served as a loading control. **E)** Quantification of thermostability of p38 MAPK western blots from four independent experiments. Table insert shows melting temperature (Tm °C) and R^2^. Error bars represent the SEM of four biological replicates. A representative image of four replicates is shown. Relative mobilities of reference proteins (masses in kilo Daltons) are shown on the left of each blot.

In order to further understand the effect of nilotinib in preventing the inflammatory response by TLR activation in human monocytes, we used *E. coli* to stimulate and induce an inflammatory response in THP-1 cells. We used live *E. coli* rather than LPS to activate multiple inflammatory pathways and not only the pathways downstream of TLR4. Similar to LPS treatment, we were able to identify the inflammatory phenotype after exposure to *E. coli* for 24h using the MALDI-TOF MS assay (**Supplementary Figure 1D** and **Supplementary Figure 5A**). Moreover, we tested by MALDI-TOF MS the p38 inhibitor losmapimod (GW856553X), which was tested in two phase III clinical trials (ClinicalTrials.gov Identifier: NCT04511819 and NCT02145468). In these results, losmapimod reduced the pro-inflammatory phenotype as well as nilotinib, but not imatinib nor MRT68601 (**Figure 4A**, **Supplementary Figure 5B** and **Supplementary Figure 1E-F**). We observed a reduction in p38α and MK2 phosphorylation in stimulated monocytes treated with nilotinib similar to the specific p38 inhibitor losmapimod (**Figure 4B** and **Supplementary Figure 5C-D**). As Nilotinib did not affect the upstream activity of MKK3, it is likely that nilotinib binds p38α directly.

**Figure 4.**
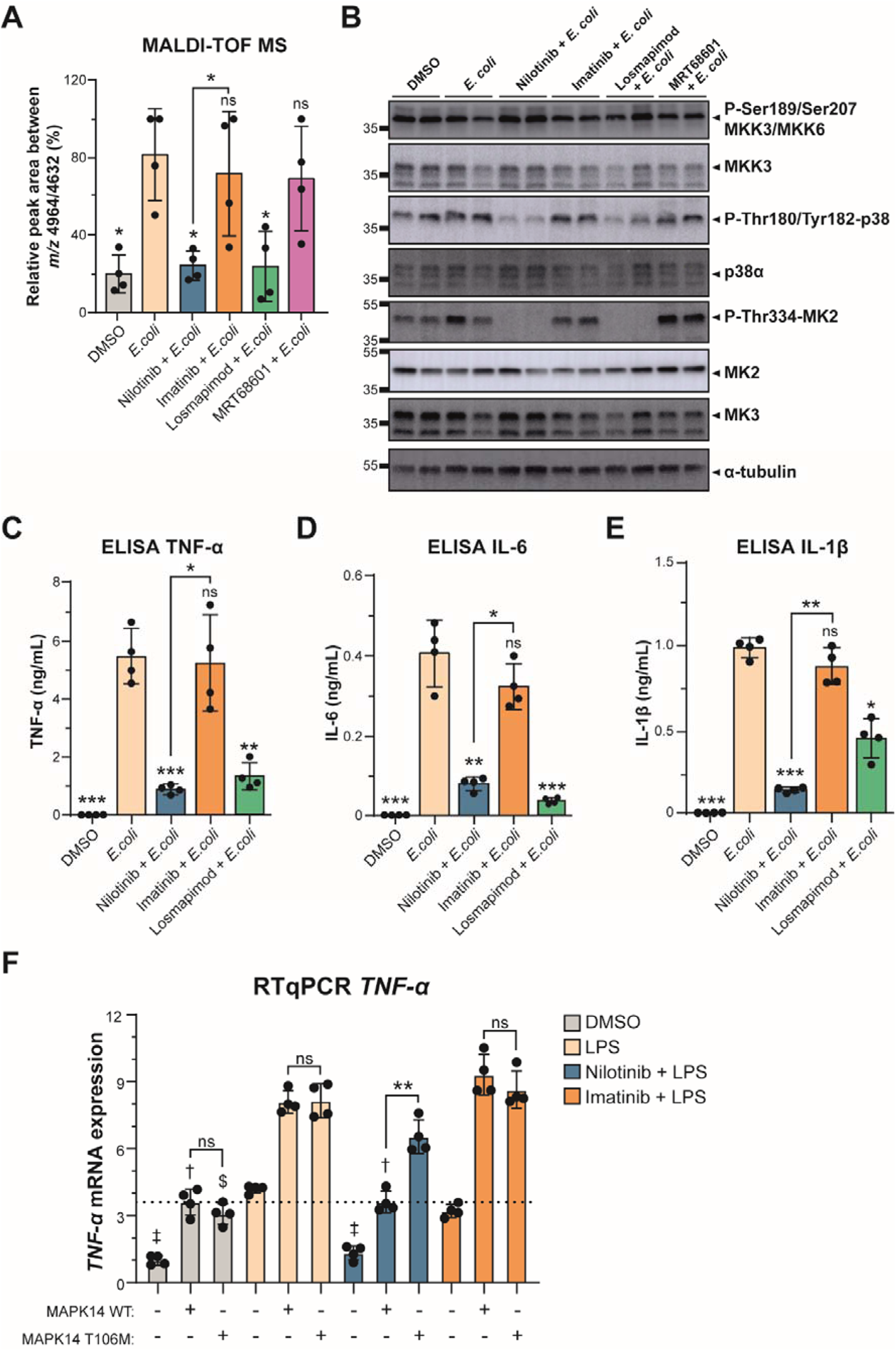
Nilotinib inhibits p38 MAPK-MK2/MK3 signalling axis. **A)** MALDI-TOF MS relative quantitation of the ratio *m/z* 4964 / 4632 in THP-1 cells pre-treated with DMSO, 5 µM nilotinib, imatinib, 1 µM losmapimod or 1 µM MRT68601 for one hour before stimulation with live *E. coli* for up to 24h. **B)** Western blot analysis of the p38 MAPK pathway in THP-1 cells pre-treated with DMSO, 5 µM nilotinib, 5 µ M imatinib, 1 µM losmapimod or 1 µM MRT68601 for one hour before stimulation with live *E. coli* for up to 15 min shows loss of p38 phosphorylation in response to nilotinib. Nilotinib and losmapimod, but not imatinib, block phosphorylation of the down-stream MK2. α-tubulin serves as a loading control. A representative image with two biological replicates of four replicates is shown. Relative mobilities of reference proteins (masses in kilo Daltons) are shown on the left of each blot. Relative quantification is shown on the Supplementary Figure 5C. **C)** TNF-α, **D)** IL-6, and **E)** IL-1β secretion was measured by ELISA at 24h. The statistical significance of the comparisons with *E. coli* is indicated as follows: ns, not significant; **, *P* ≤ 0.01; ***, *P* ≤ 0.001. **F)** *TNF-α* expression were determined by RT-qPCR in non-transfected or transfected HEK293 cells with MAPK14 Wt or MAPK14 M107T for 48h and pre-treated with DMSO, 5 µM nilotinib or imatinib for one hour before stimulation with LPS for up to 24h. The results were analyzed using the 2^−ΔΔCt^ method and normalized using *GAPDH* and *TBP* as the reference genes; and non-transfected DMSO sample as the reference sample. The statistical significance between MAPK14 Wt and MAPK14 T106M is indicated as follows: ns, not significant; ***, *P* ≤ 0.001. The statistical significance of the comparisons with LPS non-transfected is indicated as ‡; with LPS transfected with MAPK14 Wt is indicated as † with LPS transfected with MAPK14 T106M is indicated as $. Error bars represent the standard deviation of four biological replicates. Significant differences between two groups were determined by Mann-Whitney U-test.

In order to demonstrate that the suppressive effects of nilotinib on the pro-inflammatory phenotype were mediated by inhibition of the p38 MAPK pathway in THP-1 cells, we analysed the cytokine production mediated by p38 MAPK. In contrast to imatinib, nilotinib significantly reduced TNF-α, IL-6 and IL-1β levels and expression in monocytes stimulated with *E. coli* (**Figure 4C-E and Supplementary Figure 6**). We also overexpressed MAPK14 and the drug-resistant MAPK14 T106M (Eyers *et al*, 1999). We found that nilotinib was able to inhibit the effect of MAPK14, but it is not of the drug-resistant MAPK14 T106M (**Figure 4F**). Altogether, our results indicate that nilotinib targets p38 alpha, which inhibits its activity and affects the production of pro-inflammatory cytokines, thereby reducing inflammation.

### Nilotinib prevents monocyte activation

In order to better understand differences between nilotinib and imatinib treatments on monocytes, we performed quantitative proteomics of THP-1 cells pre-treated with DMSO, 5 µM nilotinib or imatinib for one hour before stimulation with live *E. coli* for up to 24h. We identified and quantified a total of 3570 proteins of which 242 proteins showed significant differences between nilotinib and imatinib treatments (**Figure 5A**, **Supplementary Table 5** and **Supplementary Figure 7A-B**). Comparing differences between nilotinib and imatinib treatments in *E. coli*-stimulated cells, we identified 25 and 30 significant proteins (log2 fold-change ≤ +/−0.6; adjusted p-value <0.05) upregulated and downregulated in nilotinib-treated cells compared to imatinib-treated cells, respectively (**Figure 5B** and **Supplementary Table 6**). We found increased levels of proteins related with actin cytoskeleton regulation, such as ROCK1 and Filamin B with nilotinib. Moreover, we found decreased levels of proteins related with cell adhesion, migration and inflammation response with nilotinib compared to imatinib or *E. coli*-stimulated cells, such as CD14, CD44, ICAM-1, IL-1β, MMP9, and TLR2 (gene ontology and pathway enrichment data in **Supplementary Table 6** and **Figure 5B-C**). During the inflammatory response, monocyte activation leads to differentiation into macrophages and also leads to an increased expression of cell adhesion proteins (Chen *et al*, 2012; Kounalakis & Corbett, 2006; Ma *et al*, 2001; Park & Kim, 2015).

**Figure 5.**
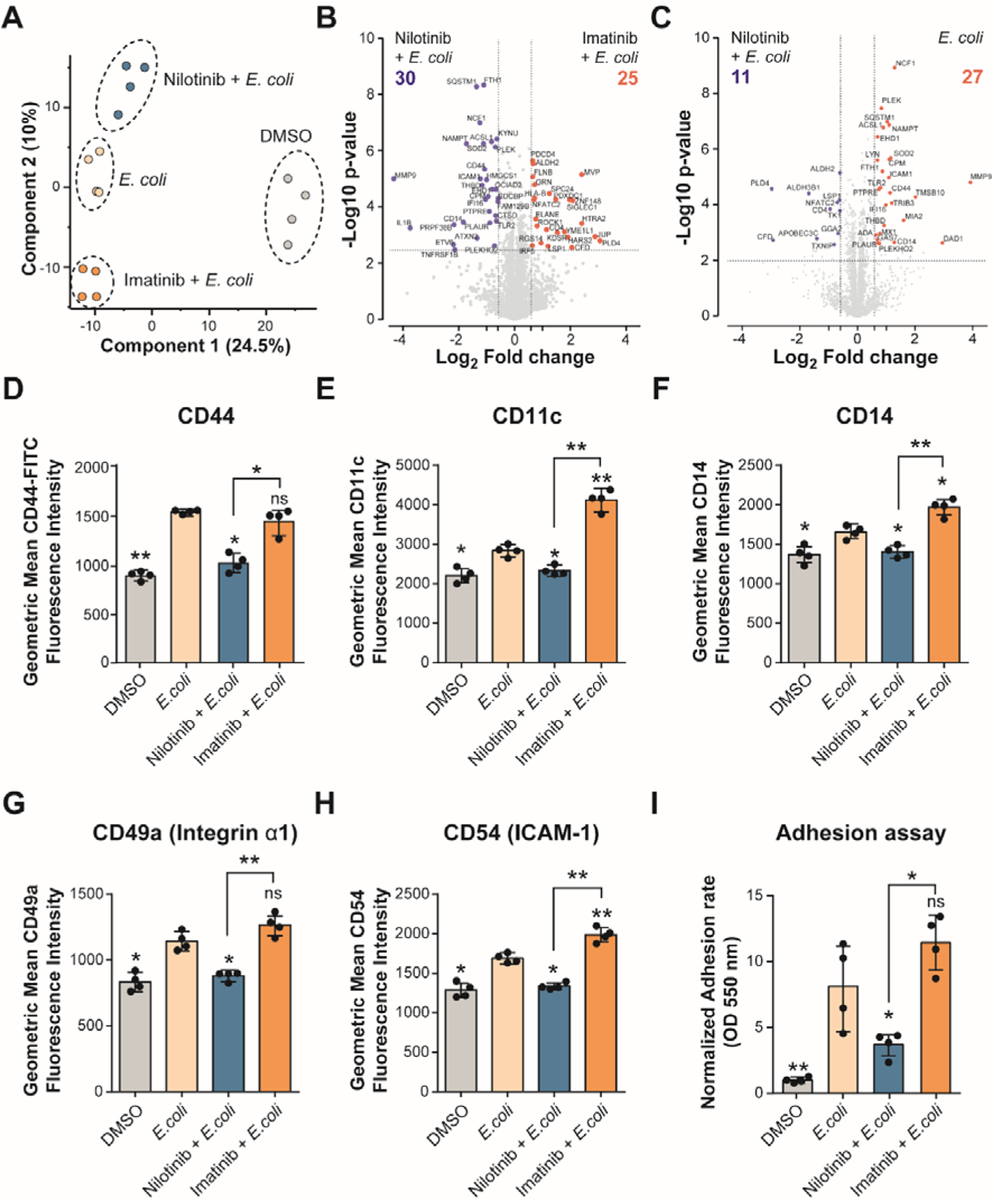
Nilotinib inhibits cell adhesion and monocyte activation. **A)** PCA plot of log2 transformed LFQ intensities from THP-1 cells pre-treated with DMSO, 5 µM nilotinib or imatinib for one hour before stimulation with live *E. coli* for up to 24h; showing distinct grouping of treatments. **B-C)** Volcano plot of THP-1 cells treated with **B)** nilotinib *vs*. imatinib, and **C)** nilotinib *vs. E. coli*, cut-off of FDR <0.05 and 1.5-fold change between conditions. **D)** Expression levels of CD44, **E)** CD11c, **F)** CD14, **G)** CD49a, and **H)** CD54 on the cell surface were measured by flow cytometry. **I)** Optical density of dissolved crystal violet was used to evaluate the adhesion rate. Significant differences between two groups were determined by Mann-Whitney U-test. The statistical significance of the comparisons with *E. coli* is indicated as follows: ns, not significant; *, *P* ≤ 0.05; **, *P* ≤ 0.01; ***, *P* ≤ 0.001. Error bars represent the standard deviation of four biological replicates.

To further understand how nilotinib affected activation of monocytes and their differentiation into macrophages upon stimulation with *E. coli,* we studied cell surface differentiation and cell adhesion markers. We validated our proteomics data by showing that MMP9 and CD44 upregulation induced by *E. coli* was abolished with nilotinib but not with imatinib (**Supplementary Figure 7C** and **Figure 5D**). Furthermore, monocyte activation markers CD11c, CD14, CD49a (integrin α1) and CD54 (ICAM-1) were suppressed with nilotinib treatment but not with imatinib (**Figure 5F-H**). Also, we observed less cell adhesion with nilotinib treatment than with imatinib (**Figure 5I**). These results indicate that nilotinib inhibits the inflammatory response of monocytes under a pro-inflammatory stimulus, inhibiting their differentiation into macrophages.

### Nilotinib reduces the inflammation phenotype in AML

Nilotinib is a second-line therapy for CML patients who failed or were intolerant to imatinib (Sasaki *et al*, 2016; Wei *et al*., 2010), but little is known about its use in other types of cancer. We performed an *in-silico* analysis of data from the Genomics of Drug Sensitivity in Cancer (GDSC) database to explore the sensitivity of nilotinib and imatinib sensitivity in different cancer types including haematological and solid tumours. We found the sensitivity to nilotinib was significantly higher than that of imatinib in haematological malignancies and gastrointestinal tumours (**Supplementary Figure 8A**). Interestingly, AML, MM, and diffuse large B cell lymphomas (DLBCL) showed significant differences between the sensitivities to nilotinib and imatinib treatments (**Supplementary Figure 8B**). These data support the interest of studying the effects of nilotinib and imatinib in myeloid malignancies as AML. In order to test whether our cellular MALDI-TOF assay can be performed in other cell lines than in THP-1 cells, and to test whether nilotinib and other TKIs can be an effective treatment in other monocyte-rich aggressive myeloid neoplasms, we used another monocytic AML cell line, OCI-AML2, and the MM cell line, NCI-H929. We also observed a significant reduction in the ratio *m/z* 4964/4632 by MALDI-TOF MS in both hematopoietic cell lines treated with nilotinib (**Supplementary Figure 8C** and **Supplementary Figure 1G-H**). Furthermore, we confirmed that nilotinib inhibits p38 MAPK phosphorylation and subsequent MK2 phosphorylation in OCI-AML2 and NCI-H929 cell lines after exposure to *E. coli* (**Supplementary Figure 8D**). Moreover, as we observed in THP-1 cells, TNF-α, IL-6, and IL-1β secretion were reduced with nilotinib treatment in OCI-AML2 (**Supplementary Figure 8E**).

In addition, we performed our MALDI-TOF assay in peripheral blood primary human monocytes and primary patient AML cells. For this, we isolated primary human CD14+ monocytes from four healthy donors and AML cells from two patients (#274 and #312). We also observed a significant reduction in the ratio m/z 4964/4632 by MALDI-TOF MS in both LPS-stimulated primary monocytes and AML samples treated with nilotinib, as well as with the third generation TKI, ponatinib, and the p38 inhibitor, losmapimod (**Figure 6A-B** and **Supplementary Figure 1I-J**), while ponatinib and bosutinib were more variable. Moreover, TNF-α secretion was reduced with nilotinib treatment in both LPS-stimulated primary monocytes and AML patient cells (**Figure 6C-D**). Taken together, these results show that our MALDI-TOF assay is suitable tool to detect pro-inflammatory phenotypes via specific biomarkers (m/z 4632 and 4964) in AML and monocytes. Furthermore, our data show that nilotinib inhibits p38 MAPK-MK2/MK3 signalling axis during the inflammatory response, reducing the levels of the pro-inflammatory cytokines in myeloma cells (**Figure 6E**).

**Figure 6.**
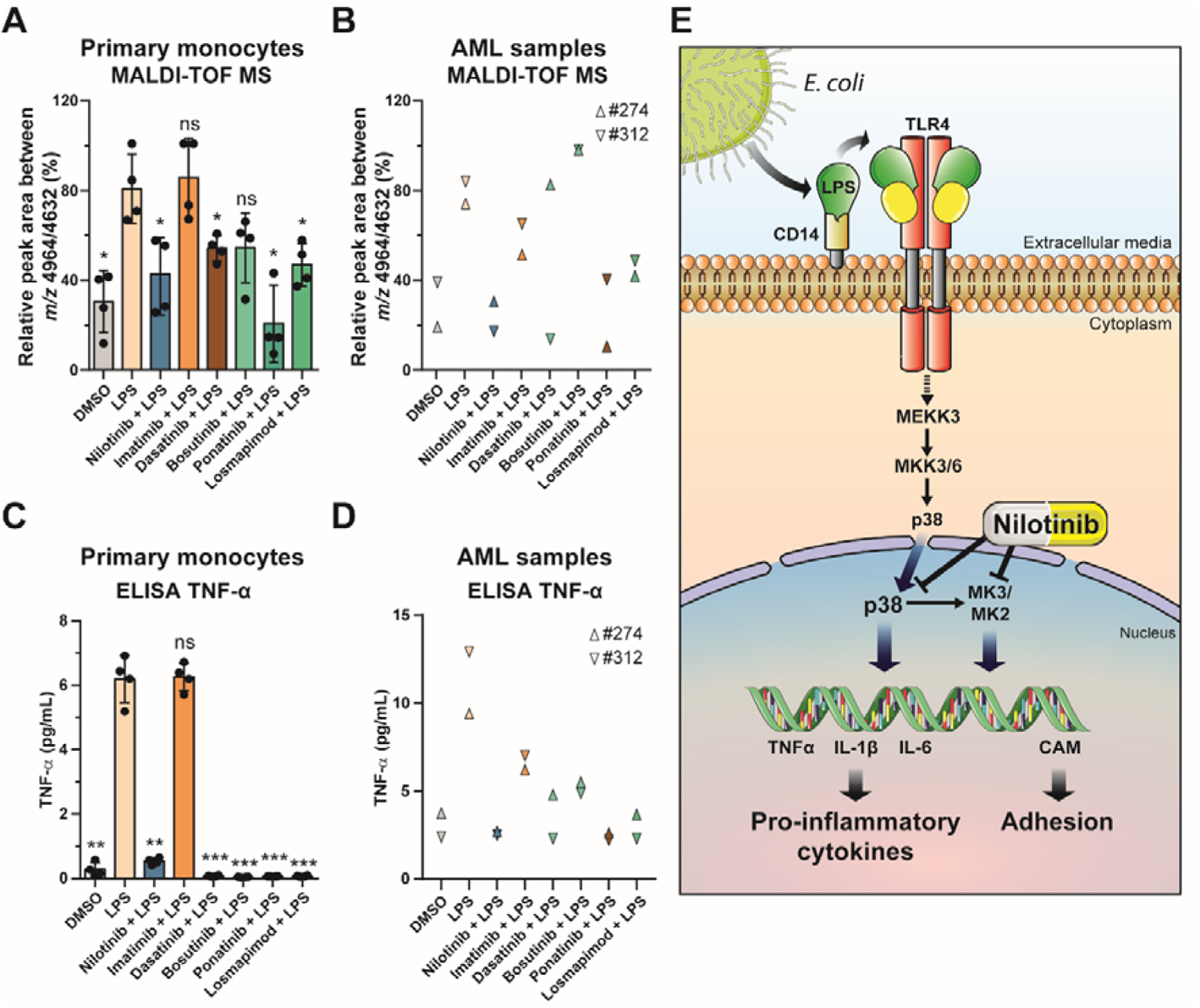
Nilotinib reduces the inflammatory phenotype in AML. **A-B)** MALDI-TOF MS relative quantitation of the ratio *m/z* 4964/4632 in **A)** human primary monocytes and **B)** primary AML cells pre-treated with DMSO, 5 µM nilotinib, imatinib, 1 µM dasatinib, bosutinib, ponatinib or losmapimod for one hour before stimulation with 100 ng/mL LPS for up to 24h. **C-D)** TNF-α secretion was measured by ELISA at 24h in **C)** human primary monocytes and **D)** primary AM cells. Error bars represent the standard deviation of four or two biological replicates. Significant differences between two groups were determined by Mann-Whitney U-test. The statistical significance of the comparisons with *E. coli* is indicated as follows: ns, not significant; *, *P* ≤ 0.05; **, *P* ≤ 0.01; ***, *P* ≤ 0.001. **E)** Effect of nilotinib during pro-inflammatory stimulus. Activation of TLR4 by a pro-inflammatory stimulus induces p38 MAPK signalling. Nilotinib inhibits p38 phosphorylation and MK2 phosphorylation reducing the levels of cytokines such as TNF-α, IL-6 or IL-1β, and cell adhesion molecules (CAMs).

## DISCUSSION

Inflammation has an important role in many aspects of acute myeloid leukaemia (AML) such as disease progression, chemoresistance, and myelosuppression (Recher, 2021). For the last five decades, AML treatment has been limited to intensive chemotherapy with cytarabine and anthracycline. Non-standard chemotherapy or immunotherapy is indicated if certain mutations or markers are detected in cancerous cells. Recently, there has been a major effort to understand the disease on the molecular level and to develop novel drugs (Kantarjian *et al*, 2021), including targeted therapies such as tyrosine kinase inhibitors (TKIs) (Fernandez *et al*, 2019). To date, 89 drugs targeting protein kinases have been clinically approved by the Food and Drug Administration (FDA, April 2021), while at least 150 are being investigated in clinical trials (Bhullar *et al*, 2018). Development of new high-throughput phenotypic drug screening approaches is important for the identification of novel drugs for the treatment of inflammation in autoimmune diseases and cancer. However, current methods for measuring pro-inflammatory cytokines are expensive as they utilize labelled antibodies, making “full deck” analyses of millions of compounds difficult (Adegbola *et al*., 2018; Neefjes & Dantuma, 2004). MALDI-TOF MS is a versatile technique with many different applications ranging from protein identification by peptide mass fingerprinting and small molecule analysis to imaging of tissues (Aichler & Walch, 2015; Cohen & Gusev, 2002). As the technology allows very fast screening, it has already found traction in the HTS field with its application to label-free screening of *in vitro* assays (De Cesare *et al*., 2018; De Cesare *et al*., 2020; Haslam *et al*, 2016; Heap *et al*., 2017; McLaren *et al*., 2021; Simon *et al*, 2021; Winter *et al*, 2018; Winter *et al*, 2019).

In this study, we present a rapid and label-free cellular MALDI-TOF MS assay able to identify an inflammatory phenotype in monocytic cells, and to screen for anti-inflammatory drugs. We applied a screen to whole cells, allowing the identification of biomarkers that provide a specific fingerprint for the phenotype of the analysed cells (Heap *et al*, 2019). Two features at *m/z* 4632 and 4964 were significantly altered upon different inflammatory stimuli, such as, LPS, Pam_2_CSK_4_, Pam_3_CSK_4_ and *E. coli* in the monocytic AML cell line THP-1 (Bosshart & Heinzelmann, 2016). The feature *m/z* 4964 has been previously described as a biomarker to identify patients with Crohn’s Disease, intestinal tuberculosis or rheumatoid arthritis using MALDI MS imaging (Kriegsmann *et al*, 2012; Zhang *et al*, 2016). It has been associated with thymosin beta 4 (TB4), a small protein involved in binding and promoting actin polymerization, however it has also been described as having anti-inflammatory properties (Holzlechner *et al*, 2017). Since the protein is only 44 amino acids, TB4 was not identified in our TPP or proteomics data and an attempt to identify it by MS/MS directly from cells by MALDI TOF MS failed. FT-ICR mass spectrometry may allow the identification of these biomarkers in the future. Moreover, we found that several pro-inflammatory markers (Cunha *et al*, 2016; Langjahr *et al*, 2014), such as TLR2, IL1B, CD14, MMP9 or SOD2 were dysregulated. We showed that the cellular MALDI-TOF MS assay can determine the inflammatory phenotype upon activation of the plasma membrane toll-like receptors (TLR), such as TLR1, TRL2, TLR4 and TLR6, but not upon activation of interferon-receptors or the intracellular TLR3 receptor.

While we tested a range of different pro-inflammatory stimuli, we found the greatest response upon LPS-stimulation, which was used in a proof-of-concept blind screen of 96 selected compounds. Within this screen, we identified nilotinib as a drug that prevents the inflammatory phenotype in stimulated AML cells by MALDI-TOF MS. However, the structurally similar drug imatinib did not have this anti-inflammatory effect. Imatinib and nilotinib are BRC-ABL tyrosine kinase inhibitors (TKIs) and the standard first-line therapy for CML (Pan *et al*., 2019). However, nilotinib possesses higher potency than imatinib against BCR-ABL and is active against most imatinib-resistant BCR-ABL mutations (Wei *et al*., 2010). The second-generation TKIs, dasatinib and bosutinib as well as the third-generation TKI, ponatinib, are also potent multitargeted tyrosine kinase inhibitors in the treatment of chronic myeloid leukemia (Jabbour *et al*, 2015). These compounds have been previously suggested to have potential anti-inflammatory effects which we confirmed in our data within a range of cell types (Akashi *et al*, 2011; Guo *et al*, 2018; Jabbour *et al*., 2015; Ozanne *et al*, 2015).

Leukemic blasts often secret inflammatory cytokines such as TNF-a, IL-1β, and IL-6, which induce expression of proteins necessary for their adhesion to vascular endothelium, migration to tissues, proliferation, and chemoresistance (Griffin *et al*, 1987; Recher, 2021). Stimulation with TLR ligands activates monocytes, which leads to the production of cytokines, chemokines, and mediators that are involved in inflammation (Stuhlmuller *et al*., 2000; Yang *et al*, 1998). TLR ligation induces the formation of a signalling complex that includes IL-1R-associated kinases (IRAKs) and TNFR-associated factor 6 (TRAF6), which mediates the K63-linked polyubiquitylation and activation of TGFβ-activated kinase 1 (TAK1). TAK1 is a MAPK kinase kinase (MAP3K) upstream of p38 MAPK and Jun N-terminal kinases (JNKs). p38 MAPK directly phosphorylates other protein kinases termed MAPK-activated protein kinases (MKs). The MKs that are phosphorylated by and functionally subordinate to p38 MAPK include MK2 and MK3, which play a versatile role in transcriptional and translational regulation, and affect inflammatory responses, triggering the production of cytokines and chemokines (Arthur & Ley, 2013; Canovas & Nebreda, 2021; Suzuki *et al*., 2018). Here, we performed a thermal proteome profiling analysis using TMTpro 16-plex (Zinn *et al*., 2021), to identify the possible different targets between nilotinib and imatinib during the inflammatory response. One of the most significant and highly reproducible targets identified was p38α as a main off-target of nilotinib, when compared to imatinib. While this technology is very powerful, one always needs to keep in mind that membrane proteins are difficult to detect with TPP and some other off-targets for nilotinib may have been missed. Nilotinib has been reported to inhibit p38α activity (Kitagawa *et al*, 2013; Li *et al*, 2011) and p38γ (Kitagawa *et al*., 2013), in an *in vitro* kinase assay and may stabilize p38α through direct interaction, as it has been observed in vascular endothelial cells and in ischemic reperfusion injury (Hadzijusufovic *et al*, 2017; Ocuin *et al*, 2012). Furthermore, we found MK3 as thermally stabilized by nilotinib, when compared to imatinib, although this could be down to changes in the phosphorylation status. However, we did not find any evidence for p38γ inhibition in our experiments, suggesting remarkable specificity for the p38α isoform. While the loss of p38 phosphorylation would suggest an inhibition upstream of p38 MAPK, we found no differences for any MAPKK and MAPKK-independent p38 phosphorylation has been described before (Beenstock *et al*, 2014; Ge *et al*, 2002; Ge *et al*, 2003). Furthermore, our results showed that nilotinib interferes with the downstream MAPK signalling pathway comparable to the specific p38 inhibitor, losmapimod (GW856553X), which has been studied in two phase III clinical trials. The potential of p38 MAPK inhibitors was initially explored within inflammatory conditions such as cancer, acute myocardial infarction, rheumatoid arthritis and Crohn’s disease, but the studies demonstrated poor clinical efficacy and unacceptable side effects (Eyers *et al*, 1998; Fisk *et al*, 2014; Lee *et al*, 1994; Lee *et al*, 2020; Wudexi *et al*, 2021). This suggests that nilotinib may be considered for additional applications outside of cancer, particularly as it is well tolerated by patients and has few side effects (Sasaki *et al*., 2016; Wei *et al*., 2010). Within the cancer field, it appears that the anti-inflammatory off-target effects from nilotinib may be beneficial as seen in tumours co-treated with doxorubicin and vincristine (Dent, 2013; Wong *et al*, 2014). Consistent with these results, nilotinib – but not imatinib – can reduce the inflammation in human monocytes-derived from AML and other myeloid cells.

Dysregulation of TLR signalling has been linked with myeloproliferative disorders (Hemmati *et al*., 2017; Monlish *et al*., 2016). While TLRs are specific for exogenous PAMPs such as lipopolysaccharide (LPS) which triggers TLR4 signalling in monocytes and macrophages, they are also activated by specific endogenous DAMPs such as HSP60 or biglycan (Takeda & Akira, 2005; Takeuchi *et al*, 1999). The bacterial surface molecule LPS activates TLR4 and CD14 in monocytes, inducing pro-inflammatory (IL-1, IL-6, and TNF-α) and then anti-inflammatory (IL-10, soluble TNF receptor, and IL-1 receptor antagonist) cytokines (Mogensen, 2009; Takeuchi *et al*., 1999). Nilotinib treatment was able to decrease the levels of IL-1β, IL-6, and TNF-α cytokines secretion upon stimulation with *E. coli* in AML cells. In line with our results, pre-incubation with nilotinib in murine macrophages derived from bone marrow led to a decreased LPS-induced *IL-6* expression (Wong *et al*, 2013). Furthermore, monocyte differentiation into macrophages induces an up-regulation of CD11c and CD14 surface markers (Chen *et al*., 2012; Ma *et al*., 2001), as well as cell adhesion molecules (Kounalakis & Corbett, 2006; Park & Kim, 2015). Interestingly, nilotinib was able to block the upregulation of CD11c, CD14, CD44, CD49a, and CD54 markers, thus reducing immune response and cell adhesion of monocytes. We further found MMP9, which cleaves CD44 and induces cell migration and cell-cell and cell-matrix adhesion (Chetty *et al*, 2012), downregulated upon nilotinib treatment. A potential mechanism of action is the Nilotinib dependent increase of ROCK1 levels in *E. coli*-stimulated cells as deficiency in ROCK1 has been implicated in the migration and recruitment of inflammatory cells such as macrophages and neutrophils during acute inflammation (Vemula *et al*, 2010). Furthermore, we observed a reduction in cell adhesion in activated monocytes upon nilotinib treatment, compared to imatinib. Our data therefore suggests that nilotinib may not only affect inflammatory signalling and immune cell adhesion, but also differentiation from monocytes into macrophages.

Our study provides a valuable tool to discover new anti-inflammatory drugs using a cellular MALDI-TOF MS assay, as we demonstrate here for nilotinib. We show that nilotinib inhibits the p38α MAPK-MK2/3 signalling axis, prevents its phosphorylation and subsequent activation, ultimately preventing the transcription of pro-inflammatory genes, cell adhesion markers and innate immunity markers. In consequence, nilotinib may have therapeutic potential through the inhibition of p38 MAPK-MK2/3 axis in inflammatory diseases as well as in myeloid malignancies such as myeloid leukaemia and multiple myeloma.

## METHODS

### Compounds

LPS was purchased from Sigma-Aldrich; Pam_2_CSK_4_, Pam_3_CSK_4_, poly(A:U) and poly(I:C) from Invitrogen; mouse IFN-γ from PeproTech; MRT68601, bosutinib, dasatinib and ponatinib from Tocris; losmapimod from Biorbyt; and nilotinib, imatinib, NG-25 and the other compounds from Supplementary Table 1 were provided by LifeArc.

### Cell culture

Acute myeloid leukaemia-derived cell lines THP-1 (ATCC) and OCI-AML2 (DSMZ), and multiple myeloma-derived cell lines NCI-H929 (ATCC) were cultured in RPMI 1640 medium (Gibco) supplemented with 10% FBS and 4 mM L-glutamine at 37°C in a humidified 5% CO_2_ atmosphere. ATCC and DSMZ routinely perform cell line authentication, using short tandem repeat profiling as a procedure. Cell experimentation was always performed within a period not exceeding six months after resuscitation in mycoplasma-free culture conditions.

### Transient transfection with lipofectamine

HEK293 (ATCC) were transfected using Lipofectamine LTX® Reagent and OptiMEM Reduced Serum Medium (Thermo Fisher) with pCMV-MAPK14 Wt (DU1765) and pCMV-MAPK14 T106M (DU3741) from MRC PPU, University of Dundee.

### Isolation of human monocytes from peripheral blood mononuclear cells (PBMC)

Mononuclear cells were freshly isolated from peripheral blood collected from healthy donor volunteers (n = 5). Blood was collected into citrate buffer. PBMC were isolated by density centrifugation using Lymphoprep (Stemcell technologies) according to manufacturer’s instructions. Monocytes were isolated using the Pan Monocyte Isolation Kit (Miltenyi Biotec) and cultured in RPMI medium supplemented with 10% FBS, 4 mM L-glutamine and 50 µg/mL penicillin/streptomycin. The study was conducted according to the principles expressed in the Helsinki Declaration and informed consent was obtained from all participants.

### AML patient cells

AML sample cells were collected after informed consent was provided via the Newcastle Haematology Biobank (Ref: 12/NE/0395). The cells were cultured in Iscove’s modified Dulbecco’s medium supplemented with 20% FBS, 4 mM L-glutamine, 50 µg/mL penicillin/streptomycin, 10 ng/mL interleukin-3 (PeproTech), and 20 ng/mL stem cell factor (PeproTech). The study was conducted according to the principles expressed in the Helsinki Declaration and informed consent was obtained from all participants. #274 sample is positive for FLT3-ITD+ and #312 sample has t(8;21) translocation.

### Blind drug screening

One million THP-1 cells/mL were incubated with each compound (**Supplementary Table 1**) at 5 µM for one hour in technical triplicates before stimulation with 100 ng/mL LPS for 24h. Compounds were staggered into 8 sets each with individual positive (NG-25) and negative controls (MRT68601) to enable maximum relative peak area calculations. This screening process was repeated in three biological replicates.

### Cell infection with *E. coli*

The DH5α strain of *E. coli* bacteria (Invitrogen) was harvested during mid-log phase. Mammalian cells were infected with a multiplicity of infection of 2. The cells with bacteria were centrifuged at 500 xg for 5 min at 37°C and incubated at 37°C in 5% CO_2_ for bacterial uptake for 15 min or 30 min. Thirty min post-infection, cells were washed once with PBS and incubated for 1h with 100 µg/mL gentamicin to kill extracellular bacteria. Then, the cells were washed with PBS twice and the media replaced with 20 µg/mL gentamicin for the remainder of the experiment.

### Cell viability assays

The viability of THP-1 cells was assessed for 24h using the Cell Proliferation Kit II (XTT) (Sigma-Aldrich) as per manufacturer’s instructions. Following incubation, the plate was read with SpectraMax iD5 microplate reader (Molecular Devices) at a wavelength of 490 nm. The reference wavelength 655 nm was also read to control for nonspecific absorption.

### Sample preparation for MALDI-TOF MS and data analysis

Cell pellets were frozen on dry ice, then thawed and washed with 100 mM Tris-HCl, pH 7.5 and centrifuged at 1,000 xg for 10 min at 4°C. 2,500 cells were spotted on target with 10 mg/mL α-cyano-4-cinnamic acid in 50% acetonitrile, 0.1% trifluoroacetic acid. Automated target spotting was performed using a Mosquito liquid handling robot (TTP Labtech). A RapifleX PharmaPulse MALDI TOF/TOF mass spectrometer (Bruker Daltonics) equipped with a Smartbeam 3D laser was used in positive ion mode. Samples were acquired in automatic mode (AutoXecute; Bruker Daltonics), totaling 10,000 shots at a 10 kHz frequency per spot. A random walk pattern (complete sample) on spot laser ablation pattern was used with an M5 Smart beam Parameter at a 45 µm × 45 µm scan range. Spot diameter was limited to 2,000 µm and a random walk pattern movement enabled at 1000 shots per raster position. Ionization was achieved using a laser power of 75% (laser attenuator offset 14%, range 30%) with a detector gain of ×6.8 in the mass range of *m/z* 2,000-20,000 with a mass suppression up to m/z 1,600. Samples were analysed in a linear geometry with optimized voltages for ion sources (ion source 1, 20 kV; pulsed ion extraction 1.3 kV), lens (8.6 kV), and a pulsed ion extraction of 180 ns. A novel 10-bit digitizer was used at a sampling rate of 1.25 GS/s. Raw data were processed first by a TopHat baseline subtraction followed by smoothing with a SavitzkyGolay algorithm. MALDI-TOF spectra were processed by FlexAnalysis Batch Process (Compass 2.0). Spectra-based PCA plots were generated using ClinPro Tools (Bruker Daltonics). A normalized relative peak area was used to measure a pro-inflammatory response between *m/z* 4632 and 4963 and was calculated with the following equation:

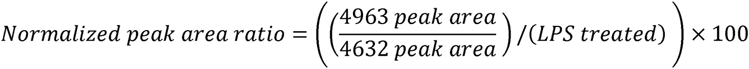

This method of intra-spectra quantification (Chen *et al*, 2017) was robust over ten different passages, where we were able to quantify the inflammatory response by MALDI-TOF MS with Z’>0.5 and *P*≤0.001.

### Quantitative RT-PCR

RNA was extracted using Trizol (Thermo Fisher Scientific). Real-time RT-qPCR from total RNA in two steps, was performed with a QuantiTect reverse transcription kit (Qiagen), and QuantiTect SYBR green kit (Qiagen) using a StepOne Applied Biosystems real-time PCR system (Thermo Fisher Scientific). Expression values of *GAPDH* and *TBP* genes in the same samples were used for normalization, using the 2^−ΔΔCT^ method. The following primers were used: *GAPDH*, forward: GTCTCCTCTGACTTCAACAGCG and, reverse: ACCACCCTGTTGCTGTAGCCAA; *IL-6*, forward: GCCCAGCTATGAACTCCTTCT and, reverse: CTTCTCCTGGGGGTACTGG; *TBP*, forward: GAGTTCCAGCGCAAGGGTTT, and, reverse: GGGTCAGTCCAGTGCCATA; *TNF-*α, forward: AAACTCATGAGCAGTCTGCA, and reverse: AGGAGATCTTCAGTTTCGGAGG.

### ELISA

Cell culture supernatant was collected at 24h post-treatment. TNF-a, IL-6, and IL-1β were measured by DuoSet ELISA kits (R&D Systems). Absorbance from four biological replicates at 450 nm was measured with the correction wavelength set at 540 nm using SpectraMax^®^ M3 microplate reader (Molecular Devices).

### Western blot

Cells were lysed using 5% SDS supplemented with Protease Inhibitor Cocktail and Phosphatase Inhibitor Cocktail 2 (Sigma-Aldrich). The following antibodies were purchased from Cell Signaling Technology (Danvers): MK2 (#3042), Thr-334-P-MK2 (#3041), MK3 (#7421), MKK3 (#8535), Ser189-P-MKK3/Ser207-P-MKK6 (#12280), p38 MAPK (#9212), p38α MAPK (#9218), p38γ MAPK (#2307) Thr180/Tyr182-P-p38 MAPK (#4511), MPP9 (#13667), anti-rabbit IgG-HRP (#7074), and anti-mouse IgG (#7076). GAPDH (sc-47724) was from Santa Cruz, and α-Tubulin (T9026) was from Sigma-Aldrich. Amersham Imager 600 digital imaging system (GE Healthcare) was used for image acquisition.

### Flow cytometry

Cell surface staining was performed by direct immunofluorescent assay with fluorescent-conjugated antibodies: CD11c-APC (#17-0114-82), CD14-AF700 (#56-0149-42), and CD54-FITC (#11-0541-82) from Thermo Fisher; CD49a-PE (#562115) and CD44-FITC (#338803) from BD Biosciences; and corresponding isotype control antibodies, for 30 min at 4°C in PBS with 1% FBS, 1% BSA and 1% human serum (Sigma) to block Fc receptors. Cells were analysed in a FACSCanto II flow cytometer (Becton-Dickinson). The results were analysed using FlowJo.

### Adhesion assay

Cells were washed twice with PBS before being fixed with 4% paraformaldehyde for 15 min. Then, cells were incubated with 5 mg/ml Crystal violet for 10 min before lysis with 2% SDS for 30 min. Absorbance from four biological replicates at 550 nm was measured using a SpectraMax^®^ M3 microplate reader (Molecular Devices). Micrographs were acquired with an inverted microscope Axio Vert.A1 FL LED (Zeiss, Cambridge).

### Cellular thermal shift assay

Cells were treated with 5 µM of nilotinib or imatinib for one hour, then 100 ng/mL LPS was added for 15 min. Cells were washed with PBS supplemented with cOmplete, EDTA-free Protease Inhibitor Cocktail (Sigma-Aldrich), 1.2 mM sodium molybdate, 1 mM sodium orthovanadate, 4 mM sodium tartrate dihydrate, 5 mM glycerophosphate, and 20 mM N-ethylmaleimide; then separated into 8 fractions for thermal profiling. Fractions were heated at 40, 44, 48, 52, 56, 60, 64 and 68°C for 3 min. For concentration compound range experiments, cells were treated with 0.1, 0.5, 1, 5 or 10 µM of nilotinib or imatinib for 1h before treatment with 100 ng/mL LPS for 15 min. Cells were heated at 56°C for 3 min. Samples were lysed with four freeze-thaw cycles using dry ice and a thermo block at 35°C. Cell lysates were centrifuged at 20,000 xg for 2h at 4°C to separate protein aggregates from soluble proteins. Supernatants were collected and used for western blot and mass spectrometry.

### Sample preparation for LC-MS/MS and label-free quantification (LFQ) data analysis

Samples were prepared for mass spectrometry using S-Trap micro columns (Protifi) according to the manufacturer’s recommended protocol. Proteins were reduced with 20 mM TCEP at 37°C for 30 min and alkylated with 20 mM N-ethylmaleimide in the dark for 30 min. A ratio of 1:10 wt:wt Trypsin TPCK Treated (Worthington-Biochem) was used to digest the samples for 2h at 47°C. Eluted peptides were dried down and resuspended in loading buffer (2% acetonitrile, 0.1% trifluoroacetic acid). Peptide samples were injected on a Dionex Ultimate 3000 RSLC (Thermo Fisher Scientific) connected to an Orbitrap Fusion Lumos Tribrid mass spectrometer (Thermo Fisher Scientific). Samples were injected on a PepMap 100 C18 LC trap column (300 µm ID x 5 mm, 5 µm, 100 Å) followed by separation on an EASY-Spray column (50 cm x 75 µm ID, PepMap C18, 2 µm, 100 Å) (Thermo Fisher Scientific). Buffer A consisted of water containing 0.1% FA and Buffer B of 80% acetonitrile containing 0.1% FA. Peptides were separated with a linear gradient of 3-35% Buffer B over 180 min followed by a step from 35-90% Buffer B in 0.5 min at 250 nL/min and held at 90% for 4 min. The gradient was then decreased to 3% Buffer B in 0.5 min at 250 nL/min, and the column equilibrated for 10 min before the next injection. Column temperature was controlled at 45°C. The Orbitrap Fusion Lumos Tribrid mass spectrometer was operated in data-dependent, positive ion mode. Full scan spectra were acquired in the range of *m/z* 400 to 1,600, at a resolution of 120,000, with an automatic gain control (AGC) target of 4×10^5^ and a maximum injection time of 50 ms. The Orbitrap was operated in ‘stop speed’ to maintain a 3 sec fixed duty cycle. The most intense precursor ions were isolated with a quadrupole mass filter width of 1.6 *m/z* and higher-energy collision-induced dissociation (HCD) fragmentation was performed in one-step collision energy of 30%. Detection of MS/MS fragments was acquired in the linear ion trap in rapid scan mode with an AGC target of 1×10^4^ ions and a maximum injection time of 45 ms. An electrospray voltage of 2.0 kV and capillary temperature of 275°C, with no sheath and auxiliary gas flow, was used.

All discovery proteomics RAW mass spectra were analysed using MaxQuant (version 1.6.10.43) (Cox & Mann, 2008) and searched against a SwissProt Homo sapiens database (containing 42,347 database entries with isoforms; downloaded on 21 April 2020). Trypsin/P was set as proteolytic enzyme, N-ethylmaleimide on cysteine was set as fixed modification, and methionine oxidation and acetylation of protein N-termini was set as variable modifications. Two missed cleavages were allowed. A protein and peptide false discovery rate (FDR) of less than 1% was employed in MaxQuant. LFQ data analysis was performed using Perseus (version 1.6.2.3) (Tyanova *et al*, 2016). Reverse hits, contaminants and proteins only identified by site were removed before downstream statistical and bioinformatics analysis. LFQ intensity data were transformed (log 2) and filtered to contain at least two unique peptides and at least three valid values in one group for comparisons. Significance testing was carried out using a two-tailed unpaired Student’s t-test and multiple hypothesis testing was controlled using the Benjamini-Hochberg FDR threshold of p-value <0.05. Gene ontology term enrichments were performed within DAVID (version 6.8) (Huang da *et al*, 2009).

### Thermal proteome profiling (TPP) and data analysis

Isobaric labelling of peptides was performed using the 16-plex tandem mass tag (TMT) reagents (Thermo Fisher Scientific) (**Supplementary Table 2**), according to the manufacturer recommended protocol. According to the different temperature points, labelled peptides were combined and desalted with C18 Macro Spin Columns (Harvard Apparatus, Holliston, MA, USA). The TMT-labelled samples were subjected to fractionation using basic-pH reversed-phase liquid chromatography on a Dionex Ultimate 3000 HPLC system (Thermo Fisher Scientific) using a Gemini C18 column (250 mm × 3 mm, 3 μm, 110 Å; Phenomenex). Buffer A consisted of 20 mM ammonium formate pH 8.0 and Buffer B of 100% acetonitrile. Peptides were fractionated using a linear gradient of 1-49% buffer B over 49 min at 250 nl/min. Twelve fractions were dried under vacuum centrifugation and resuspended in 2% acetonitrile with 0.1% TFA for LC-MS/MS analysis. Labelled peptide samples were injected on a Dionex Ultimate 3000 RSLC (Thermo Fisher Scientific) connected to an Orbitrap Fusion Lumos Tribrid mass spectrometer (Thermo Fisher Scientific). Buffer A consisted of water containing 0.1% FA and Buffer B of 80% acetonitrile containing 0.1% FA. Peptides were separated with a linear gradient of 3-35% Buffer B over 180 min followed by a step from 35-90% Buffer B in 0.5 min at 250 nL/min and held at 90% for 4 min. The gradient was then decreased to 3% Buffer B in 0.5 min at 250 nL/min, and the column equilibrated for 10 min before the next injection. The scan sequence began with an MS1 spectrum (Orbitrap analysis; resolution 120,000; mass range *m/z* 375-1500; automatic gain control (AGC) target 4x10E5; maximum injection time 50 ms). MS2 analysis consisted of collision-induced dissociation (CID); AGC 1x10E4; normalized collision energy (NCE) of 30%; maximum injection time 50 ms; and isolation window of 0.7. Following acquisition of each MS2 spectrum, MS3 precursors were fragmented by high energy collision-induced dissociation (HCD) and analysed using the Orbitrap (NCE 55%; AGC 5x10E4; maximum injection time 86 ms, resolution was 50,000) and synchronous precursor selection (SPS) was enabled to include 10 MS/MS fragment ions in the FTMS3 scan.

Mass spectrometry data analysis was performed similarly as described above, but TMT modification on the peptide N-termini or lysine residues were enabled for the 16-plex TMT reagents, and deamidation of asparagine and glutamine was included as variable modification in MaxQuant. TPP data analysis was performed using the TPP packages (https://www.bioconductor.org/packages/release/bioc/html/TPP.html) in the R statistical programming language. The data were normalized as detailed in Miettinen *et al*. (Miettinen *et al*., 2018) (**Supplementary Table 3**). Proteins that changed significantly in the four replicates between nilotinib and imatinib with a standard deviation below two degrees in the melting temperature were selected as positive hits.

### Data availability

The mass spectrometry proteomics data have been deposited to the ProteomeXchange Consortium (Deutsch *et al*, 2017) via the PRIDE partner repository (Perez-Riverol *et al*, 2019) with the data set identifier: PXD025020.

Reviewer account details: Username: Password: H0T2rCOL

### Statistical analysis

The Shapiro-Wilk test was used to check expression data sets for normality, and the Levene test was used for homogeneity of variances. Mann-Whitney U-test was performed in all the analysis, excluding proteomics and thermal proteome profiling.

## ACKNOWLEDGEMENTS

We would like to thank Abeer Dannoura, Wezi Sendama, Marie-Hélène Ruchaud-Sparagano, Akshada Gajbhiye, Julien Peltier, and Anetta Härtlova for their technical support; and María Villa-Morales for a critical reading of this manuscript. This research was partly funded by a Wellcome Trust Investigator Award (215542/Z/19/Z) and generous start-up funding of Newcastle University to M.T. R.E.H. was funded by an iCASE studentship by the BBSRC and Bruker Daltonics. M.E.D. is a Marie Sklodowska Curie Fellow within the European Union’s Horizon 2020 research and innovation programme under the Marie Skłodowska-Curie grant agreement No. 890296. For the purpose of Open Access, the author has applied a CC BY public copyright license to any Author Accepted Manuscript version arising from this submission. Professor Simpson is a National Institute for Health Research (NIHR) Senior Investigator. The views expressed in this article are those of the authors and not necessarily those of the NIHR, or the Department of Health and Social Care.

## AUTHORSHIP CONTRIBUTIONS

J.L.M.-R. and R.E.H. are equal co-first authors. R.E.H. developed the MALDI-TOF MS assay. J.L.M.-R. and R.E.H. performed most of the experiments. J.L.M.-R. and T.H. performed LFQ and TMT mass spectrometry-based proteomics. T.H. analysed TPP experiments. M.E.D. performed and analysed additional MALDI-TOF MS experiments. J.L.M.-R. and J.I. performed and analysed RT qPCR assays. B.S. provided blind drug panel. J.S. and A.J.S. provided PMBCs from healthy donors. J.M.A., H.B. and O.H. provided AML patient samples. M.T. and J.L.M-R conceived the original idea and provided supervision. J.L.M.-R., R.E.H., and M.T. wrote the manuscript. All authors provided critical feedback and helped shape the research, analysis, and manuscript.

## SUPPLEMENTAL INFORMATION

The online version of this article contains supplementary data.

## REFERENCES

Adegbola SO, Sahnan K, Warusavitarne J, Hart A, Tozer P (2018) Anti-TNF Therapy in Crohn’s Disease. Int J Mol Sci 19

Aichler M, Walch A (2015) MALDI Imaging mass spectrometry: current frontiers and perspectives in pathology research and practice. Lab Invest 95: 422–431

Akashi N, Matsumoto I, Tanaka Y, Inoue A, Yamamoto K, Umeda N, Tanaka Y, Hayashi T, Goto D, Ito S et al (2011) Comparative suppressive effects of tyrosine kinase inhibitors imatinib and nilotinib in models of autoimmune arthritis. Mod Rheumatol 21: 267–275

Arthur JS, Ley SC (2013) Mitogen-activated protein kinases in innate immunity. Nat Rev Immunol 13: 679–692

Becher I, Werner T, Doce C, Zaal EA, Togel I, Khan CA, Rueger A, Muelbaier M, Salzer E, Berkers CR et al (2016) Thermal profiling reveals phenylalanine hydroxylase as an off-target of panobinostat. Nat Chem Biol 12: 908–910

Beenstock J, Ben-Yehuda S, Melamed D, Admon A, Livnah O, Ahn NG, Engelberg D (2014) The p38beta mitogen-activated protein kinase possesses an intrinsic autophosphorylation activity, generated by a short region composed of the alpha-G helix and MAPK insert. J Biol Chem 289: 23546–23556

Bhullar KS, Lagaron NO, McGowan EM, Parmar I, Jha A, Hubbard BP, Rupasinghe HPV (2018) Kinase-targeted cancer therapies: progress, challenges and future directions. Mol Cancer 17: 48

Binder S, Luciano M, Horejs-Hoeck J (2018) The cytokine network in acute myeloid leukemia (AML): A focus on pro- and anti-inflammatory mediators. Cytokine Growth Factor Rev 43: 8–15

Bosshart H, Heinzelmann M (2016) THP-1 cells as a model for human monocytes. Ann Transl Med 4: 438

Butcher EC (2005) Can cell systems biology rescue drug discovery? Nat Rev Drug Discov 4: 461–467

Canovas B, Nebreda AR (2021) Diversity and versatility of p38 kinase signalling in health and disease. Nat Rev Mol Cell Biol 22: 346–366

Chandler J, Haslam C, Hardy N, Leveridge M, Marshall P (2017) A Systematic Investigation of the Best Buffers for Use in Screening by MALDI-Mass Spectrometry. SLAS Discov 22: 1262–1269

Chen RF, Wang L, Cheng JT, Yang KD (2012) Induction of IFNalpha or IL-12 depends on differentiation of THP-1 cells in dengue infections without and with antibody enhancement. BMC Infect Dis 12: 340

Chen X, Wo F, Chen J, Tan J, Wang T, Liang X, Wu J (2017) Ratiometric Mass Spectrometry for Cell Identification and Quantitation Using Intracellular “Dual-Biomarkers”. Sci Rep 7: 17432

Chetty C, Vanamala SK, Gondi CS, Dinh DH, Gujrati M, Rao JS (2012) MMP-9 induces CD44 cleavage and CD44 mediated cell migration in glioblastoma xenograft cells. Cell Signal 24: 549–559

Cohen LH, Gusev AI (2002) Small molecule analysis by MALDI mass spectrometry. Anal Bioanal Chem 373: 571–586

Cox J, Mann M (2008) MaxQuant enables high peptide identification rates, individualized p.p.b.-range mass accuracies and proteome-wide protein quantification. Nat Biotechnol 26: 1367–1372

Craver BM, El Alaoui K, Scherber RM, Fleischman AG (2018) The Critical Role of Inflammation in the Pathogenesis and Progression of Myeloid Malignancies. Cancers (Basel) 10

Cunha C, Gomes C, Vaz AR, Brites D (2016) Exploring New Inflammatory Biomarkers and Pathways during LPS-Induced M1 Polarization. Mediators Inflamm 2016: 6986175

De Cesare V, Johnson C, Barlow V, Hastie J, Knebel A, Trost M (2018) The MALDI-TOF E2/E3 Ligase Assay as Universal Tool for Drug Discovery in the Ubiquitin Pathway. Cell Chem Biol 25: 1117–1127 e1114

De Cesare V, Moran J, Traynor R, Knebel A, Ritorto MS, Trost M, McLauchlan H, Hastie CJ, Davies P (2020) High-throughput matrix-assisted laser desorption/ionization time-of-flight (MALDI-TOF) mass spectrometry-based deubiquitylating enzyme assay for drug discovery. Nat Protoc 15: 4034–4057

Dent P (2013) The flip side of doxorubicin: Inflammatory and tumor promoting cytokines. Cancer Biol Ther 14: 774–775

Deutsch EW, Csordas A, Sun Z, Jarnuczak A, Perez-Riverol Y, Ternent T, Campbell DS, Bernal-Llinares M, Okuda S, Kawano S et al (2017) The ProteomeXchange consortium in 2017: supporting the cultural change in proteomics public data deposition. Nucleic Acids Res 45: D1100–D1106

Dreisewerd K (2014) Recent methodological advances in MALDI mass spectrometry. Anal Bioanal Chem 406: 2261–2278

Dulai PS, Sandborn WJ (2016) Next-Generation Therapeutics for Inflammatory Bowel Disease. Curr Gastroenterol Rep 18: 51

Eyers PA, Craxton M, Morrice N, Cohen P, Goedert M (1998) Conversion of SB 203580-insensitive MAP kinase family members to drug-sensitive forms by a single amino-acid substitution. Chem Biol 5: 321–328

Eyers PA, van den IP, Quinlan RA, Goedert M, Cohen P (1999) Use of a drug-resistant mutant of stress-activated protein kinase 2a/p38 to validate the in vivo specificity of SB 203580. FEBS Lett 451: 191–196

Fernandez S, Desplat V, Villacreces A, Guitart AV, Milpied N, Pigneux A, Vigon I, Pasquet JM, Dumas PY (2019) Targeting Tyrosine Kinases in Acute Myeloid Leukemia: Why, Who and How? Int J Mol Sci 20

Fisk M, Gajendragadkar PR, Maki-Petaja KM, Wilkinson IB, Cheriyan J (2014) Therapeutic potential of p38 MAP kinase inhibition in the management of cardiovascular disease. Am J Cardiovasc Drugs 14: 155–165

Franken H, Mathieson T, Childs D, Sweetman GM, Werner T, Togel I, Doce C, Gade S, Bantscheff M, Drewes G et al (2015) Thermal proteome profiling for unbiased identification of direct and indirect drug targets using multiplexed quantitative mass spectrometry. Nat Protoc 10: 1567–1593

Ge B, Gram H, Di Padova F, Huang B, New L, Ulevitch RJ, Luo Y, Han J (2002) MAPKK-independent activation of p38alpha mediated by TAB1-dependent autophosphorylation of p38alpha. Science 295: 1291–1294

Ge B, Xiong X, Jing Q, Mosley JL, Filose A, Bian D, Huang S, Han J (2003) TAB1beta (transforming growth factor-beta-activated protein kinase 1-binding protein 1beta), a novel splicing variant of TAB1 that interacts with p38alpha but not TAK1. J Biol Chem 278: 2286–2293

Gerets HH, Dhalluin S, Atienzar FA (2011) Multiplexing cell viability assays. Methods Mol Biol 740: 91–101

Gordon S, Taylor PR (2005) Monocyte and macrophage heterogeneity. Nat Rev Immunol 5: 953–964

Griffin JD, Rambaldi A, Vellenga E, Young DC, Ostapovicz D, Cannistra SA (1987) Secretion of interleukin-1 by acute myeloblastic leukemia cells in vitro induces endothelial cells to secrete colony stimulating factors. Blood 70: 1218–1221

Guitot K, Drujon T, Burlina F, Sagan S, Beaupierre S, Pamlard O, Dodd RH, Guillou C, Bolbach G, Sachon E et al (2017) A direct label-free MALDI-TOF mass spectrometry based assay for the characterization of inhibitors of protein lysine methyltransferases. Anal Bioanal Chem 409: 3767–3777

Guo K, Bu X, Yang C, Cao X, Bian H, Zhu Q, Zhu J, Zhang D (2018) Treatment Effects of the Second-Generation Tyrosine Kinase Inhibitor Dasatinib on Autoimmune Arthritis. Front Immunol 9: 3133

Hadzijusufovic E, Albrecht-Schgoer K, Huber K, Hoermann G, Grebien F, Eisenwort G, Schgoer W, Herndlhofer S, Kaun C, Theurl M et al (2017) Nilotinib-induced vasculopathy: identification of vascular endothelial cells as a primary target site. Leukemia 31: 2388–2397

Haslam C, Hellicar J, Dunn A, Fuetterer A, Hardy N, Marshall P, Paape R, Pemberton M, Resemannand A, Leveridge M (2016) The Evolution of MALDI-TOF Mass Spectrometry toward Ultra-High-Throughput Screening: 1536-Well Format and Beyond. J Biomol Screen 21: 176–186

Heap RE, Hope AG, Pearson LA, Reyskens K, McElroy SP, Hastie CJ, Porter DW, Arthur JSC, Gray DW, Trost M (2017) Identifying Inhibitors of Inflammation: A Novel High-Throughput MALDI-TOF Screening Assay for Salt-Inducible Kinases (SIKs). SLAS Discov 22: 1193–1202

Heap RE, Segarra-Fas A, Blain AP, Findlay GM, Trost M (2019) Profiling embryonic stem cell differentiation by MALDI TOF mass spectrometry: development of a reproducible and robust sample preparation workflow. Analyst 144: 6371–6381

Hemmati S, Haque T, Gritsman K (2017) Inflammatory Signaling Pathways in Preleukemic and Leukemic Stem Cells. Front Oncol 7: 265

Holzlechner M, Strasser K, Zareva E, Steinhauser L, Birnleitner H, Beer A, Bergmann M, Oehler R, Marchetti-Deschmann M (2017) In Situ Characterization of Tissue-Resident Immune Cells by MALDI Mass Spectrometry Imaging. J Proteome Res 16: 65–76

Huang da W, Sherman BT, Lempicki RA (2009) Systematic and integrative analysis of large gene lists using DAVID bioinformatics resources. Nat Protoc 4: 44–57

Inglese J, Johnson RL, Simeonov A, Xia M, Zheng W, Austin CP, Auld DS (2007) High-throughput screening assays for the identification of chemical probes. Nat Chem Biol 3: 466–479

Jabbour E, Kantarjian H, Cortes J (2015) Use of second- and third-generation tyrosine kinase inhibitors in the treatment of chronic myeloid leukemia: an evolving treatment paradigm. Clin Lymphoma Myeloma Leuk 15: 323–334

Jarzab A, Kurzawa N, Hopf T, Moerch M, Zecha J, Leijten N, Bian Y, Musiol E, Maschberger M, Stoehr G et al (2020) Meltome atlas-thermal proteome stability across the tree of life. Nat Methods 17: 495–503

Kantarjian H, Kadia T, DiNardo C, Daver N, Borthakur G, Jabbour E, Garcia-Manero G, Konopleva M, Ravandi F (2021) Acute myeloid leukemia: current progress and future directions. Blood Cancer J 11: 41

Kim EK, Choi EJ (2010) Pathological roles of MAPK signaling pathways in human diseases. Biochim Biophys Acta 1802: 396–405

Kitagawa D, Yokota K, Gouda M, Narumi Y, Ohmoto H, Nishiwaki E, Akita K, Kirii Y (2013) Activity-based kinase profiling of approved tyrosine kinase inhibitors. Genes Cells 18: 110–122

Kounalakis NS, Corbett SA (2006) Lipopolysaccharide transiently activates THP-1 cell adhesion. J Surg Res 135: 137–143

Kriegsmann M, Seeley EH, Schwarting A, Kriegsmann J, Otto M, Thabe H, Dierkes B, Biehl C, Sack U, Wellmann A et al (2012) MALDI MS imaging as a powerful tool for investigating synovial tissue. Scand J Rheumatol 41: 305–309

Langjahr P, Diaz-Jimenez D, De la Fuente M, Rubio E, Golenbock D, Bronfman FC, Quera R, Gonzalez MJ, Hermoso MA (2014) Metalloproteinase-dependent TLR2 ectodomain shedding is involved in soluble toll-like receptor 2 (sTLR2) production. PLoS One 9: e104624

Lee JC, Laydon JT, McDonnell PC, Gallagher TF, Kumar S, Green D, McNulty D, Blumenthal MJ, Heys JR, Landvatter SW et al (1994) A protein kinase involved in the regulation of inflammatory cytokine biosynthesis. Nature 372: 739–746

Lee S, Rauch J, Kolch W (2020) Targeting MAPK Signaling in Cancer: Mechanisms of Drug Resistance and Sensitivity. Int J Mol Sci 21

Li P, Zheng Y, Chen X (2017) Drugs for Autoimmune Inflammatory Diseases: From Small Molecule Compounds to Anti-TNF Biologics. Front Pharmacol 8: 460

Li YY, An J, Jones SJ (2011) A computational approach to finding novel targets for existing drugs. PLoS Comput Biol 7: e1002139

Ma W, Lim W, Gee K, Aucoin S, Nandan D, Kozlowski M, Diaz-Mitoma F, Kumar A (2001) The p38 mitogen-activated kinase pathway regulates the human interleukin-10 promoter via the activation of Sp1 transcription factor in lipopolysaccharide-stimulated human macrophages. J Biol Chem 276: 13664–13674

Malik N, Vollmer S, Nanda SK, Lopez-Pelaez M, Prescott A, Gray N, Cohen P (2015) Suppression of interferon beta gene transcription by inhibitors of bromodomain and extra-terminal (BET) family members. Biochem J 468: 363–372

Mateus A, Kurzawa N, Becher I, Sridharan S, Helm D, Stein F, Typas A, Savitski MM (2020) Thermal proteome profiling for interrogating protein interactions. Mol Syst Biol 16: e9232

Mateus A, Maatta TA, Savitski MM (2016) Thermal proteome profiling: unbiased assessment of protein state through heat-induced stability changes. Proteome Sci 15: 13

McLaren DG, Shah V, Wisniewski T, Ghislain L, Liu C, Zhang H, Saldanha SA (2021) High-Throughput Mass Spectrometry for Hit Identification: Current Landscape and Future Perspectives. SLAS Discov 26: 168–191

Michelini E, Cevenini L, Mezzanotte L, Coppa A, Roda A (2010) Cell-based assays: fuelling drug discovery. Anal Bioanal Chem 398: 227–238

Miettinen TP, Peltier J, Hartlova A, Gierlinski M, Jansen VM, Trost M, Bjorklund M (2018) Thermal proteome profiling of breast cancer cells reveals proteasomal activation by CDK4/6 inhibitor palbociclib. EMBO J 37

Mogensen TH (2009) Pathogen recognition and inflammatory signaling in innate immune defenses. Clin Microbiol Rev 22: 240–273, Table of Contents

Monlish DA, Bhatt ST, Schuettpelz LG (2016) The Role of Toll-Like Receptors in Hematopoietic Malignancies. Front Immunol 7: 390

Neefjes J, Dantuma NP (2004) Fluorescent probes for proteolysis: tools for drug discovery. Nat Rev Drug Discov 3: 58–69

Newman AC, Scholefield CL, Kemp AJ, Newman M, McIver EG, Kamal A, Wilkinson S (2012) TBK1 kinase addiction in lung cancer cells is mediated via autophagy of Tax1bp1/Ndp52 and non-canonical NF-kappaB signalling. PLoS One 7: e50672

Ocuin LM, Zeng S, Cavnar MJ, Sorenson EC, Bamboat ZM, Greer JB, Kim TS, Popow R, DeMatteo RP (2012) Nilotinib protects the murine liver from ischemia/reperfusion injury. J Hepatol 57: 766–773

Ozanne J, Prescott AR, Clark K (2015) The clinically approved drugs dasatinib and bosutinib induce anti-inflammatory macrophages by inhibiting the salt-inducible kinases. Biochem J 465: 271–279

Pan P, Wang L, Wang Y, Shen L, Zheng P, Bi C, Zhang A, Lv Y, Xue Z, Sun M et al (2019) Systematic Review and Meta-Analysis of - New-Generation Tyrosine Kinase Inhibitors versus Imatinib for Newly Diagnosed Chronic Myeloid Leukemia. Acta Haematol: 1–13

Park GS, Kim JH (2015) LPS Up-Regulates ICAM-1 Expression in Breast Cancer Cells by Stimulating a MyD88-BLT2-ERK-Linked Cascade, Which Promotes Adhesion to Monocytes. Mol Cells 38: 821–828

Perez-Riverol Y, Csordas A, Bai J, Bernal-Llinares M, Hewapathirana S, Kundu DJ, Inuganti A, Griss J, Mayer G, Eisenacher M et al (2019) The PRIDE database and related tools and resources in 2019: improving support for quantification data. Nucleic Acids Res 47: D442–D450

Pophali PA, Patnaik MM (2016) The Role of New Tyrosine Kinase Inhibitors in Chronic Myeloid Leukemia. Cancer J 22: 40–50

Recher C (2021) Clinical Implications of Inflammation in Acute Myeloid Leukemia. Front Oncol 11: 623952

Reinhard FB, Eberhard D, Werner T, Franken H, Childs D, Doce C, Savitski MF, Huber W, Bantscheff M, Savitski MM et al (2015) Thermal proteome profiling monitors ligand interactions with cellular membrane proteins. Nat Methods 12: 1129–1131

Ritorto MS, Ewan R, Perez-Oliva AB, Knebel A, Buhrlage SJ, Wightman M, Kelly SM, Wood NT, Virdee S, Gray NS et al (2014) Screening of DUB activity and specificity by MALDI-TOF mass spectrometry. Nat Commun 5: 4763

Saei AA, Beusch CM, Sabatier P, Wells JA, Gharibi H, Meng Z, Chernobrovkin A, Rodin S, Nareoja K, Thorsell AG et al (2021) System-wide identification and prioritization of enzyme substrates by thermal analysis. Nat Commun 12: 1296

Sasaki K, Kantarjian H, Jabbour E, Ravandi F, Takahashi K, Konopleva M, Borthakur G, Garcia-Manero G, Wierda W, Daver N et al (2016) Frontline therapy with high-dose imatinib versus second generation tyrosine kinase inhibitor in patients with chronic-phase chronic myeloid leukemia - a propensity score analysis. Haematologica 101: e324–327

Savitski MM, Reinhard FB, Franken H, Werner T, Savitski MF, Eberhard D, Martinez Molina D, Jafari R, Dovega RB, Klaeger S et al (2014) Tracking cancer drugs in living cells by thermal profiling of the proteome. Science 346: 1255784

Simon RP, Winter M, Kleiner C, Wehrle L, Karnath M, Ries R, Zeeb M, Schnapp G, Fiegen D, Habe TT et al (2021) MALDI-TOF-Based Affinity Selection Mass Spectrometry for Automated Screening of Protein-Ligand Interactions at High Throughput. SLAS Discov 26: 44–57

Smith MJ, Ivanov DP, Weber RJM, Wingfield J, Viant MR (2021) Acoustic Mist Ionization Mass Spectrometry for Ultrahigh-Throughput Metabolomics Screening. Anal Chem 93: 9258–9266

Stuhlmuller B, Ungethum U, Scholze S, Martinez L, Backhaus M, Kraetsch HG, Kinne RW, Burmester GR (2000) Identification of known and novel genes in activated monocytes from patients with rheumatoid arthritis. Arthritis Rheum 43: 775–790

Suzuki T, Sakata K, Mizuno N, Palikhe S, Yamashita S, Hattori K, Matsuda N, Hattori Y (2018) Different involvement of the MAPK family in inflammatory regulation in human pulmonary microvascular endothelial cells stimulated with LPS and IFN-gamma. Immunobiology 223: 777–785

Takeda K, Akira S (2005) Toll-like receptors in innate immunity. Int Immunol 17: 1–14

Takeuchi O, Hoshino K, Kawai T, Sanjo H, Takada H, Ogawa T, Takeda K, Akira S (1999) Differential roles of TLR2 and TLR4 in recognition of gram-negative and gram-positive bacterial cell wall components. Immunity 11: 443–451

Tyanova S, Temu T, Sinitcyn P, Carlson A, Hein MY, Geiger T, Mann M, Cox J (2016) The Perseus computational platform for comprehensive analysis of (prote)omics data. Nat Methods 13: 731–740

Vemula S, Shi J, Hanneman P, Wei L, Kapur R (2010) ROCK1 functions as a suppressor of inflammatory cell migration by regulating PTEN phosphorylation and stability. Blood 115: 1785–1796

Wang KY, Chuang SA, Lin PC, Huang LS, Chen SH, Ouarda S, Pan WH, Lee PY, Lin CC, Chen YJ (2008) Multiplexed immunoassay: quantitation and profiling of serum biomarkers using magnetic nanoprobes and MALDI-TOF MS. Anal Chem 80: 6159–6167

Wei G, Rafiyath S, Liu D (2010) First-line treatment for chronic myeloid leukemia: dasatinib, nilotinib, or imatinib. J Hematol Oncol 3: 47

Winter M, Bretschneider T, Kleiner C, Ries R, Hehn JP, Redemann N, Luippold AH, Bischoff D, Buttner FH (2018) Establishing MALDI-TOF as Versatile Drug Discovery Readout to Dissect the PTP1B Enzymatic Reaction. SLAS Discov 23: 561–573

Winter M, Ries R, Kleiner C, Bischoff D, Luippold AH, Bretschneider T, Buttner FH (2019) Automated MALDI Target Preparation Concept: Providing Ultra-High-Throughput Mass Spectrometry-Based Screening for Drug Discovery. SLAS Technol 24: 209–221

Wong J, Smith LB, Magun EA, Engstrom T, Kelley-Howard K, Jandhyala DM, Thorpe CM, Magun BE, Wood LJ (2013) Small molecule kinase inhibitors block the ZAK-dependent inflammatory effects of doxorubicin. Cancer Biol Ther 14: 56–63

Wong J, Tran LT, Magun EA, Magun BE, Wood LJ (2014) Production of IL-1beta by bone marrow-derived macrophages in response to chemotherapeutic drugs: synergistic effects of doxorubicin and vincristine. Cancer Biol Ther 15: 1395–1403

Wudexi I, Shokri E, Abo-Aly M, Shindo K, Abdel-Latif A (2021) Comparative Effectiveness of Anti-Inflammatory Drug Treatments in Coronary Heart Disease Patients: A Systematic Review and Network Meta-Analysis. Mediators Inflamm 2021: 5160728

Xu W, Mo J, Ocak U, Travis ZD, Enkhjargal B, Zhang T, Wu P, Peng J, Li T, Zuo Y et al (2020) Activation of Melanocortin 1 Receptor Attenuates Early Brain Injury in a Rat Model of Subarachnoid Hemorrhage viathe Suppression of Neuroinflammation through AMPK/TBK1/NF-kappaB Pathway in Rats. Neurotherapeutics 17: 294–308

Yang RB, Mark MR, Gray A, Huang A, Xie MH, Zhang M, Goddard A, Wood WI, Gurney AL, Godowski PJ (1998) Toll-like receptor-2 mediates lipopolysaccharide-induced cellular signalling. Nature 395: 284–288

Zhang F, Xu C, Ning L, Hu F, Shan G, Chen H, Yang M, Chen W, Yu J, Xu G (2016) Exploration of Serum Proteomic Profiling and Diagnostic Model That Differentiate Crohn’s Disease and Intestinal Tuberculosis. PLoS One 11: e0167109

Zinn N, Werner T, Doce C, Mathieson T, Boecker C, Sweetman G, Fufezan C, Bantscheff M (2021) Improved Proteomics-Based Drug Mechanism-of-Action Studies Using 16-Plex Isobaric Mass Tags. J Proteome Res 20: 1792–1801

