## Supplementary Figures for "A MALDI-TOF assay identifies nilotinib as an inhibitor of inflammation in acute myeloid leukaemia"

#### Supplementary Figure 1

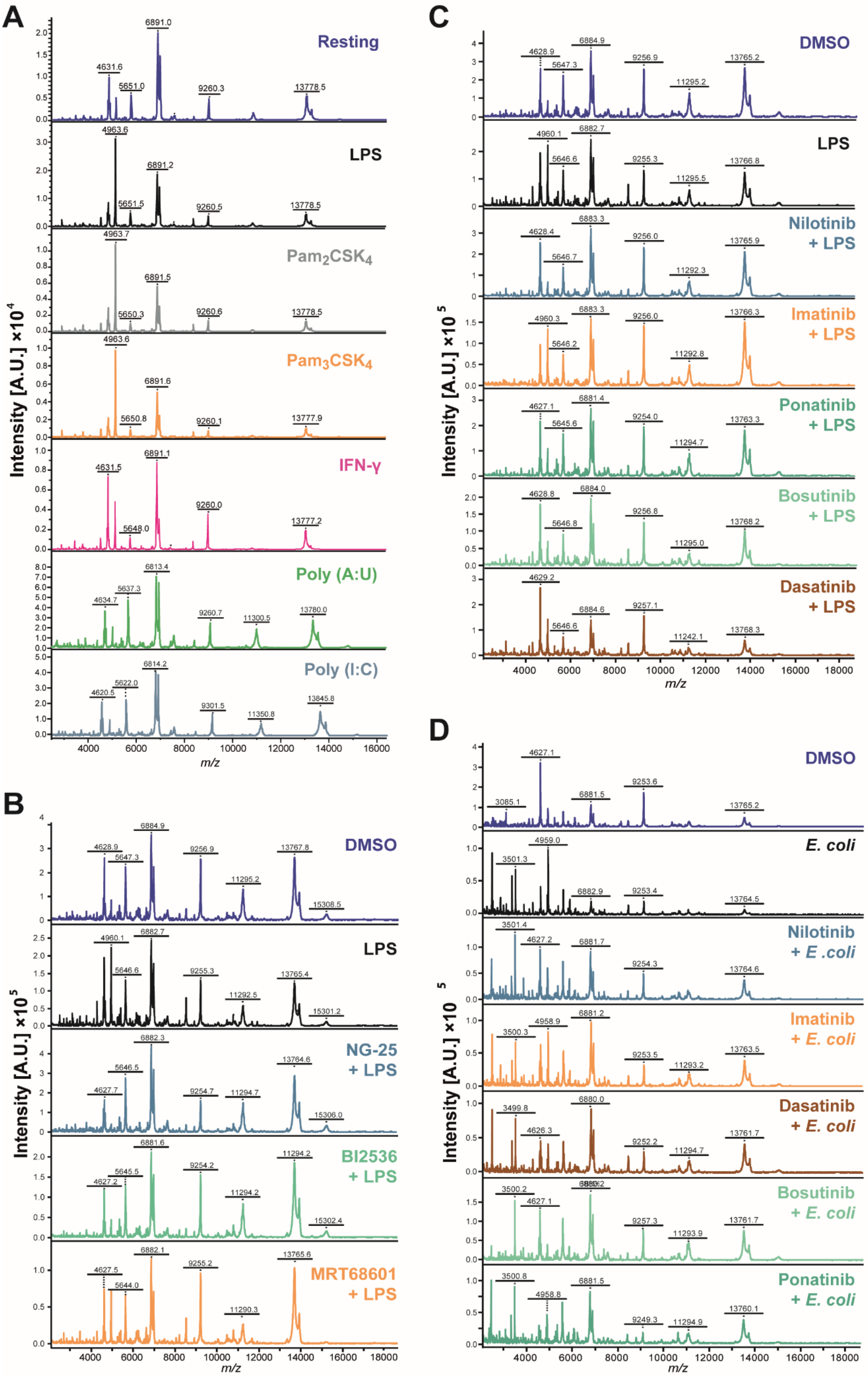

#### Supplementary Figure 1

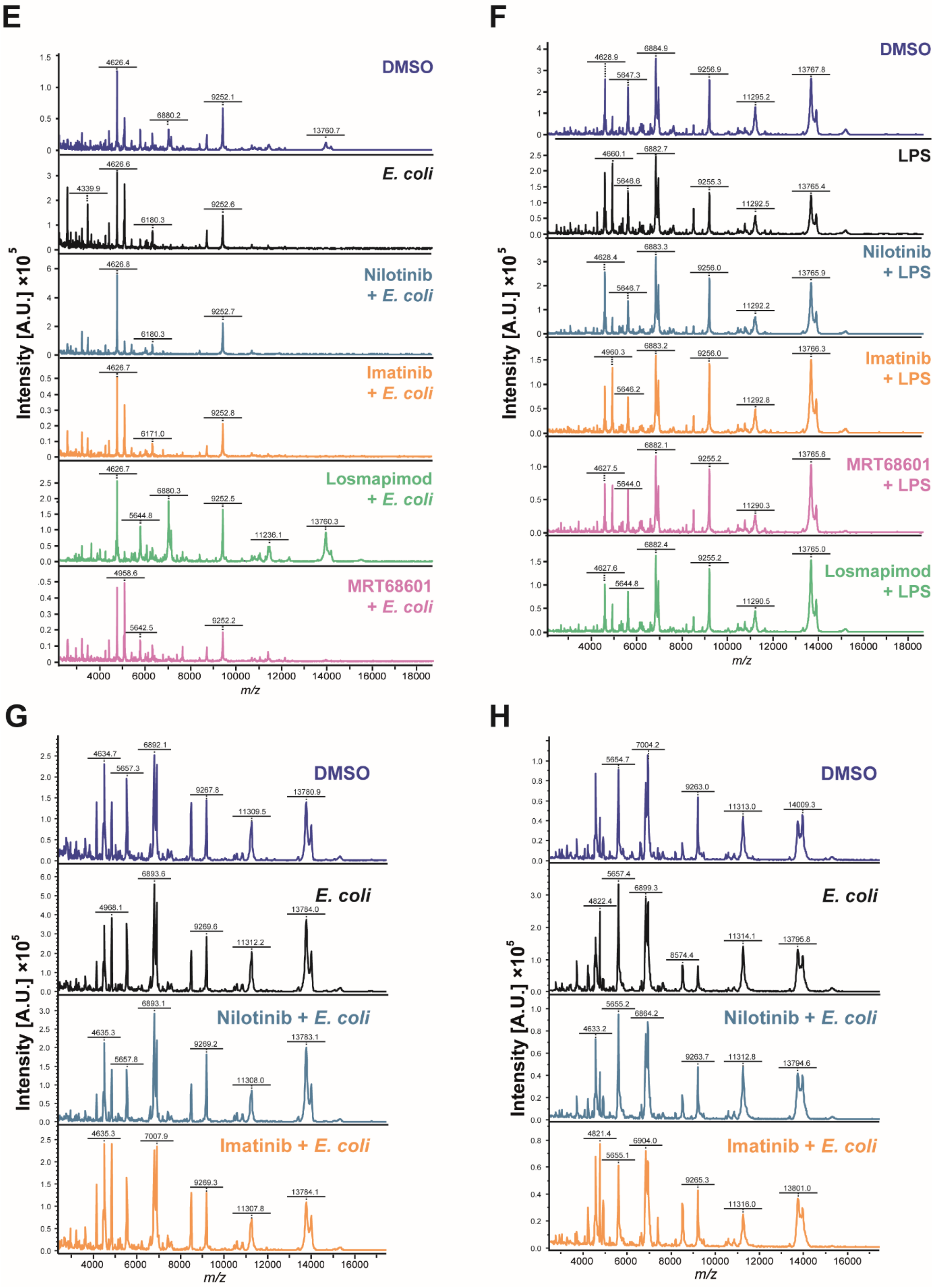

Figure EV1

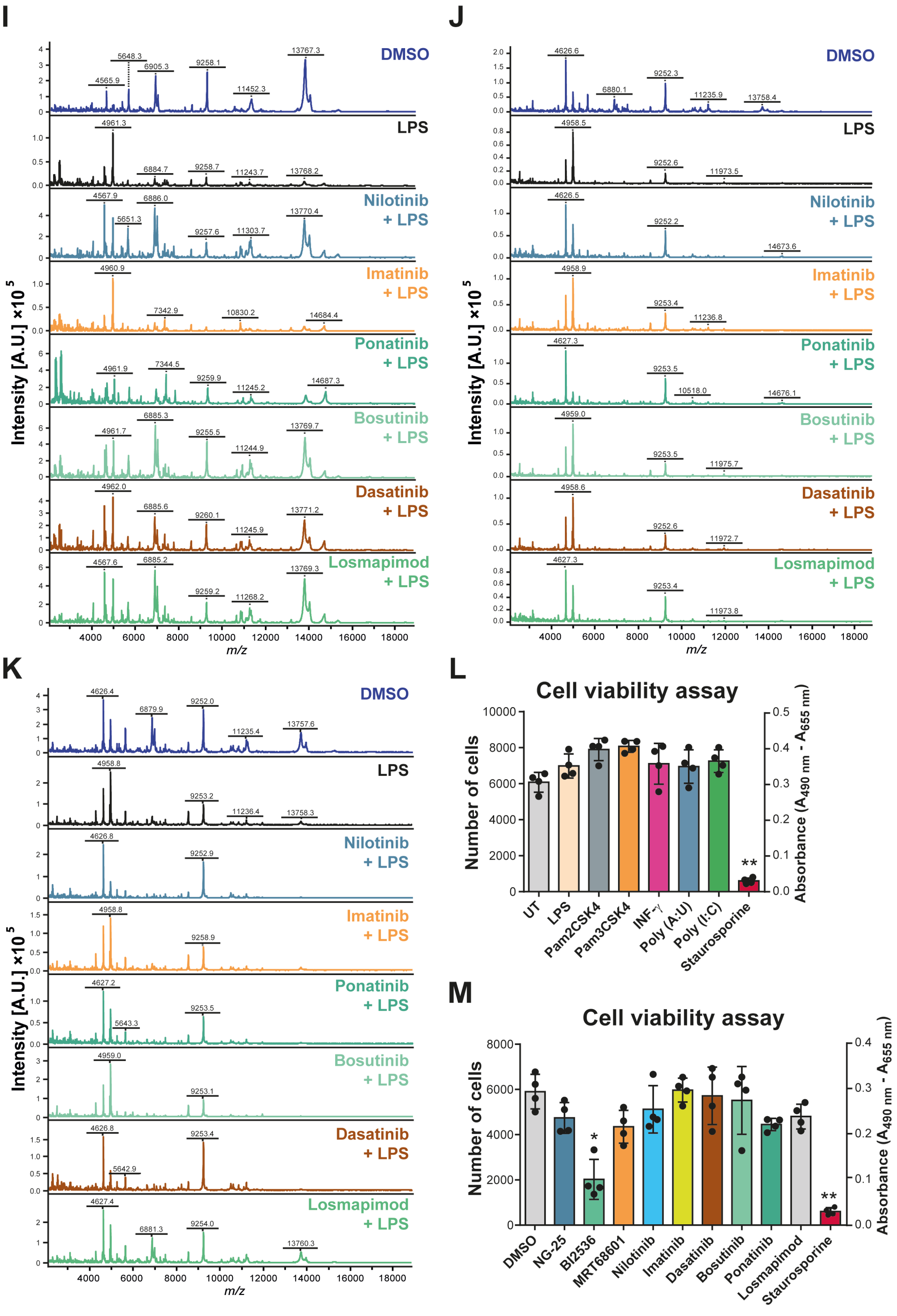
I
J
K
L
M
DMSO
LPS
Nilotinib + LPS
Imatinib + LPS
Ponatinib + LPS
Bosutinib + LPS
Dasatinib + LPS
Losmapimod + LPS
Intensity [A.U.] ×10^5
m/z
Cell viability assay
Number of cells
Absorbance (A490 nm - A655 nm)
UT
LPS
Pam2CSK4
Pam3CSK4
INF-γ
Poly (A:U)
Poly (I:C)
Staurosporine
DMSO
NG-25
BI2536
MRT68601
Nilotinib
Imatinib
Dasatinib
Bosutinib
Ponatinib
Losmapimod
Staurosporine

**Supplementary Figure 1. MALDI-TOF MS spectra. A)** Representative MALDI-TOF MS spectra of untreated (resting) THP-1 cells and treated with 100 ng/mL LPS, 100 ng/mL $Pam_2CSK_4$, 100 ng/mL $Pam_3CSK_4$, 100 U/mL IFN-γ, 1 μg/mL poly(I:C) or 1 μg/mL poly(A:U) for 24h. **B)** Representative MALDI-TOF MS spectra of THP-1 cells pre-treated with vehicle control (DMSO), 5 μM NG-25, 5 μM BI2536 or 1 μM MRT68601 for one hour before 100 ng/mL LPS-treatment for up to 24h. **C-D)** Representative MALDI-TOF MS spectra of THP-1 cells pre-treated with DMSO, 5 μM nilotinib, 5 μM imatinib, 1 μM dasatinib, 1 μM bosutinib or 1 μM ponatinib C) for one hour before 100 ng/mL LPS-treatment for up to 24h; D) for one hour before stimulation with live *E. coli* for up to 24h. **E-F)** Representative MALDI-TOF MS spectra of THP-1 cells pre-treated with DMSO, 5 μM nilotinib, 5 μM imatinib, 1 μM losmapimod or 1 μM MRT68601 E) for one hour before stimulation with live *E. coli* for up to 24h; F) for one hour before 100 ng/mL LPS-treatment for up to 24h. **G-H)** Representative MALDI-TOF MS spectra of G) OCI-AML2 and H) NCI-H929 cells pre-treated with DMSO, 5 μM nilotinib or imatinib for one hour before stimulation with live *E. coli* for up to 24h. **I-K)** Representative MALDI-TOF MS spectra of I) primary monocytes and J-K) primary AML cells pre-treated with DMSO, 5 μM nilotinib, 5 μM imatinib, 1 μM dasatinib, 1 μM bosutinib, 1 μM ponatinib or 1 μM losmapimod for one hour before 100 ng/mL LPS-treatment for up to 24h. **L** and **M**) Represent XTT cell viability assays. Significant differences between two groups were determined by Mann-Whitney U-test. The statistical significance of the comparisons with resting is indicated as follows: **, $P \leq 0.01$; *, $P \leq 0.05$. Error bars represent the standard deviation of four biological replicates.

#### Supplementary Figure 2

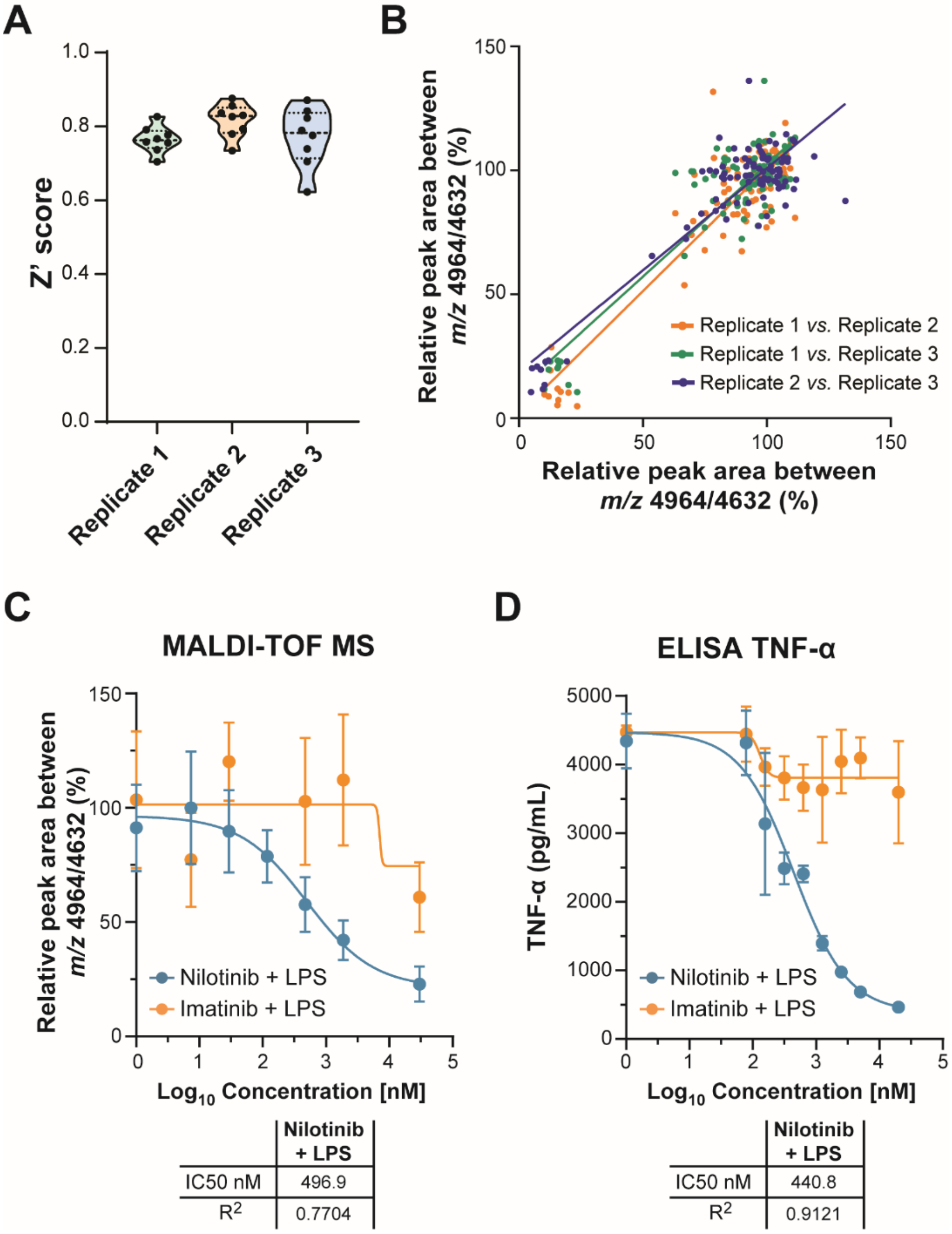


**Supplementary Figure 2. MALDI-TOF MS screening quality control. A**) Z' scores of biological replicates 1, 2 and 3 from the 96 compounds screened across the 8 staggered sets. **B)** Correlation plots of biological replicates 1, 2 and 3 showing correlation $R^2 > 0.80$ and $P \leq 0.001$. **C)** MALDI-TOF MS IC50 curve of nilotinib- and imatinib-pre-treated cells one hour before 100 ng/mL LPS-treatment for up to 24h. **D)** TNF-α secretion IC50 curve of nilotinib- and imatinib-treated THP-1 cells. Table below shows IC50 concentration (nM) and $R^2$. Error bars represent the SEM of six (**C**) or four (**D**) biological replicates.

### Supplementary Figure 3

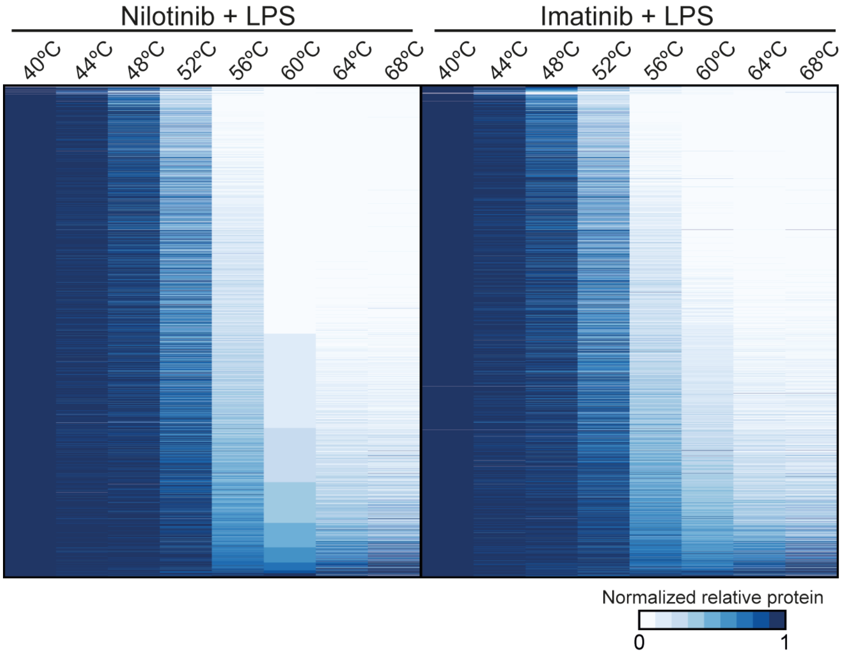


**Supplementary Figure 3. TPP ratio heatmap.** Heat map of protein thermal stabilities in THP-1 cells with 5 µM nilotinib and imatinib for one hour before 100 ng/mL LPS-treatment for up to 15 min. The median relative abundance across replicates (n = 4) at the indicated temperature is shown for each protein as fold change relative to the lowest measured temperature (40 °C).

**Supplementary Figure 4**

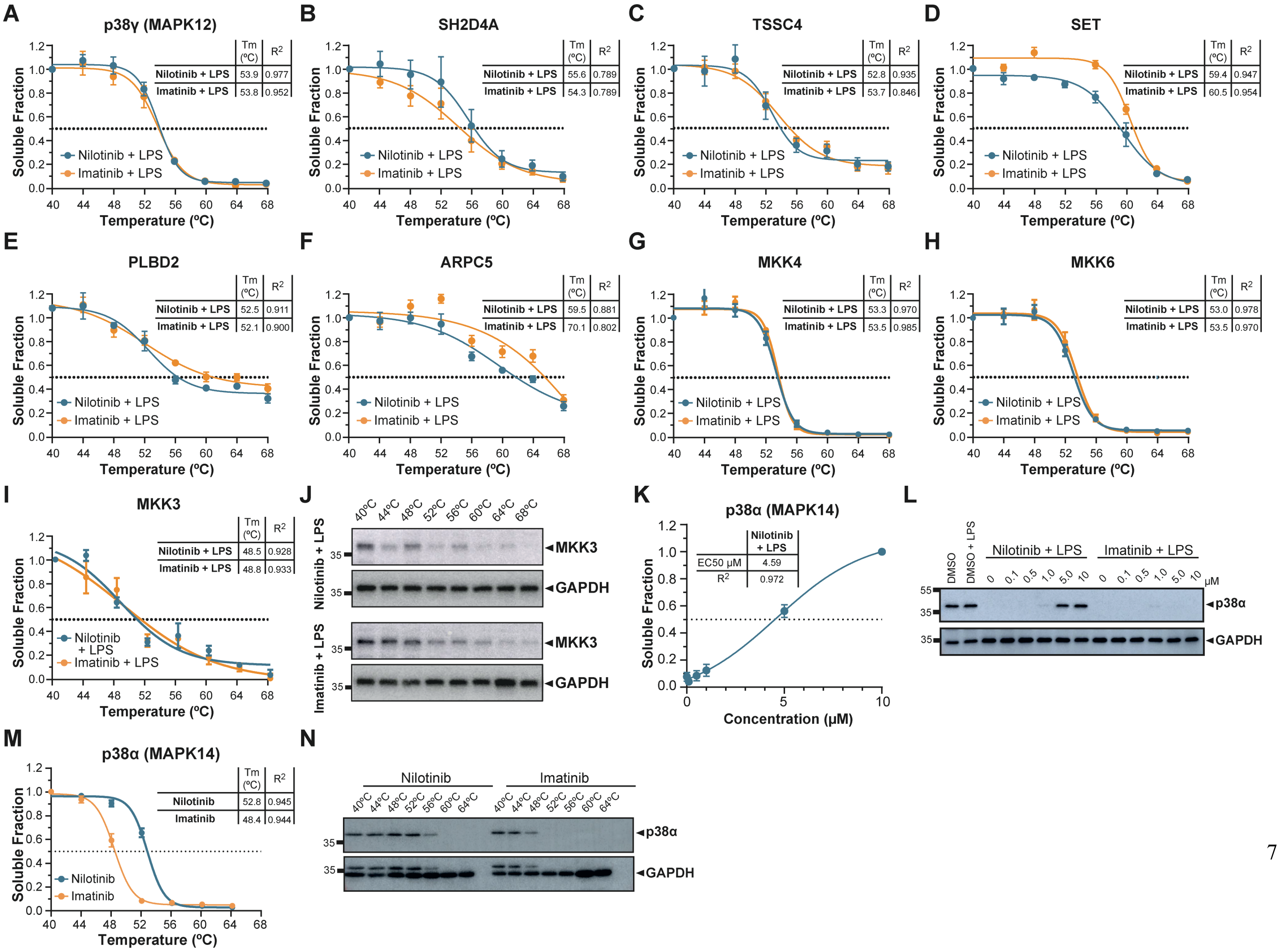

**Supplementary Figure 4. TPP analysis. A)** Determination by TPP analysis of the thermostability of p38 gamma (MAPK12), **B)** SH2 domain-containing protein 4A (SH2D4A), **C)** protein TSSC4, **D)** protein SET, **E)** putative phospholipase B-like 2 (PLBD2), **F)** Actin-related protein 2/3 complex subunit 5 (ARPC5), **G)** Mitogen-activated protein kinase kinase 4 (MKK4), **H)** Mitogen-activated protein kinase kinase 6 (MKK6) protein at the indicated temperatures with 5 µM nilotinib or imatinib for one hour before 100 ng/mL LPS-treatment for up to 15 min. Table insert shows melting temperature (Tm ºC) and $R^2$. **I-J)** Determination of the thermostability of MKK3 at the indicated temperatures in THP-1 cells pre-treated with 5 µM nilotinib or imatinib for one hour before 100 ng/mL LPS-treatment for up to 15 min. J) Representative western blots of the thermostability of MKK3. GAPDH served as a loading control. Table insert shows melting temperature (Tm ºC) and $R^2$. **K-L)** Determination of the thermostability of p38 alpha (MAPK12) at 56ºC in DMSO, 100 ng/mL LPS and pre-treated with the indicated concentrations of nilotinib or imatinib for one hour before 100 ng/mL LPS-treatment for up to 15 min. Table insert below shows effective concentration (EC50) (µM) and $R^2$. **M-N)** Determination of the thermostability of p38 alpha (MAPK12) at 56ºC in 5 µM nilotinib or imatinib for one hour. Table insert shows melting temperature (Tm ºC) and $R^2$. Error bars represent the SEM of four biological replicates. GAPDH served as a loading control. A representative image of four replicates is shown. Relative mobilities of reference proteins (masses in kilo Daltons) are shown on the left of each blot.

#### Supplementary Figure 5

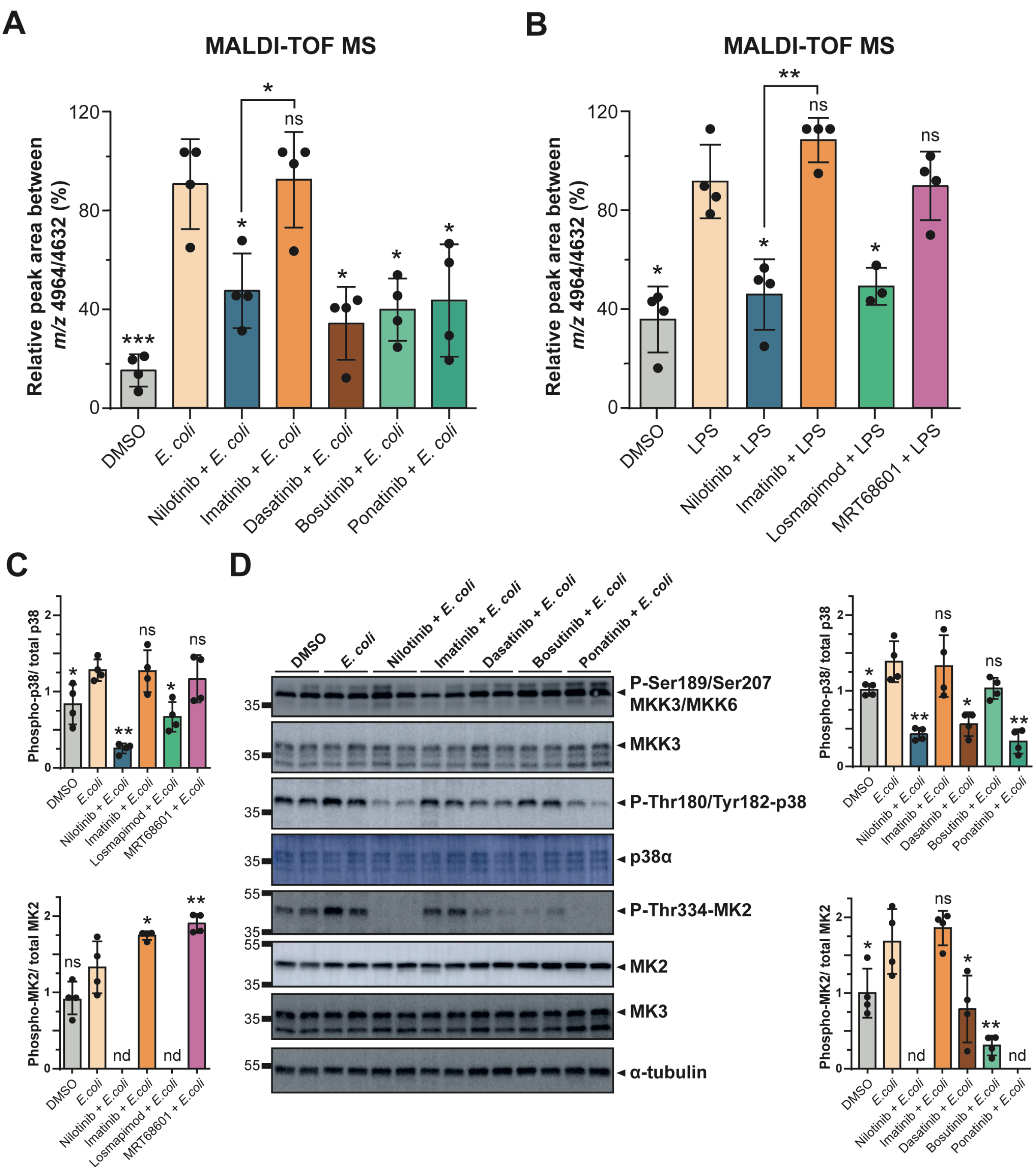


**Supplementary Figure 5. Nilotinib reduces the pro-inflammatory phenotype. A)** MALDI-TOF MS relative quantitation of the ratio *m/z* 4964 / 4632 in THP-1 cells pre-treated with DMSO, 5 µM nilotinib, imatinib, 1 µM dasatinib, bosutinib or ponatinib for one hour before stimulation with live *E. coli* for up to 24h. **B)** MALDI-TOF MS relative quantitation of the ratio *m/z* 4964 / 4632 in THP-1 cells pre-treated with DMSO, 5 µM nilotinib, imatinib, 1 µM losmapimod or MRT68601 for one hour before stimulation with LPS for up to 24h. **C)** Relative quantification of four replicates (Figure 4B) is shown on the right for p38 and MK2 phosphorylation levels. **D)** Western blot analysis of the p38 MAPK pathway in THP-1 cells pre-treated with DMSO, 5 µM nilotinib, 5 µM nilotinib, imatinib, 1 µM dasatinib, bosutinib or ponatinib for one hour before stimulation with live *E. coli* for up to 15 min. α-tubulin serves as a loading control. A representative image with two biological replicates of four replicates is shown. Relative mobilities of reference proteins (masses in kilo Daltons) are shown on the left of each blot. Relative quantification is shown on the right for p38 and MK2 phosphorylation levels. Error bars represent the standard deviation of four biological replicates. Significant differences between two groups were determined by a Mann-Whitney U-test. The statistical significance of the comparisons with *E. coli* is indicated as follows: ns, not significant; nd, not detected; *, $P \leq 0.05$; **, $P \leq 0.01$; ***, $P \leq 0.001$.

#### Supplementary Figure 6

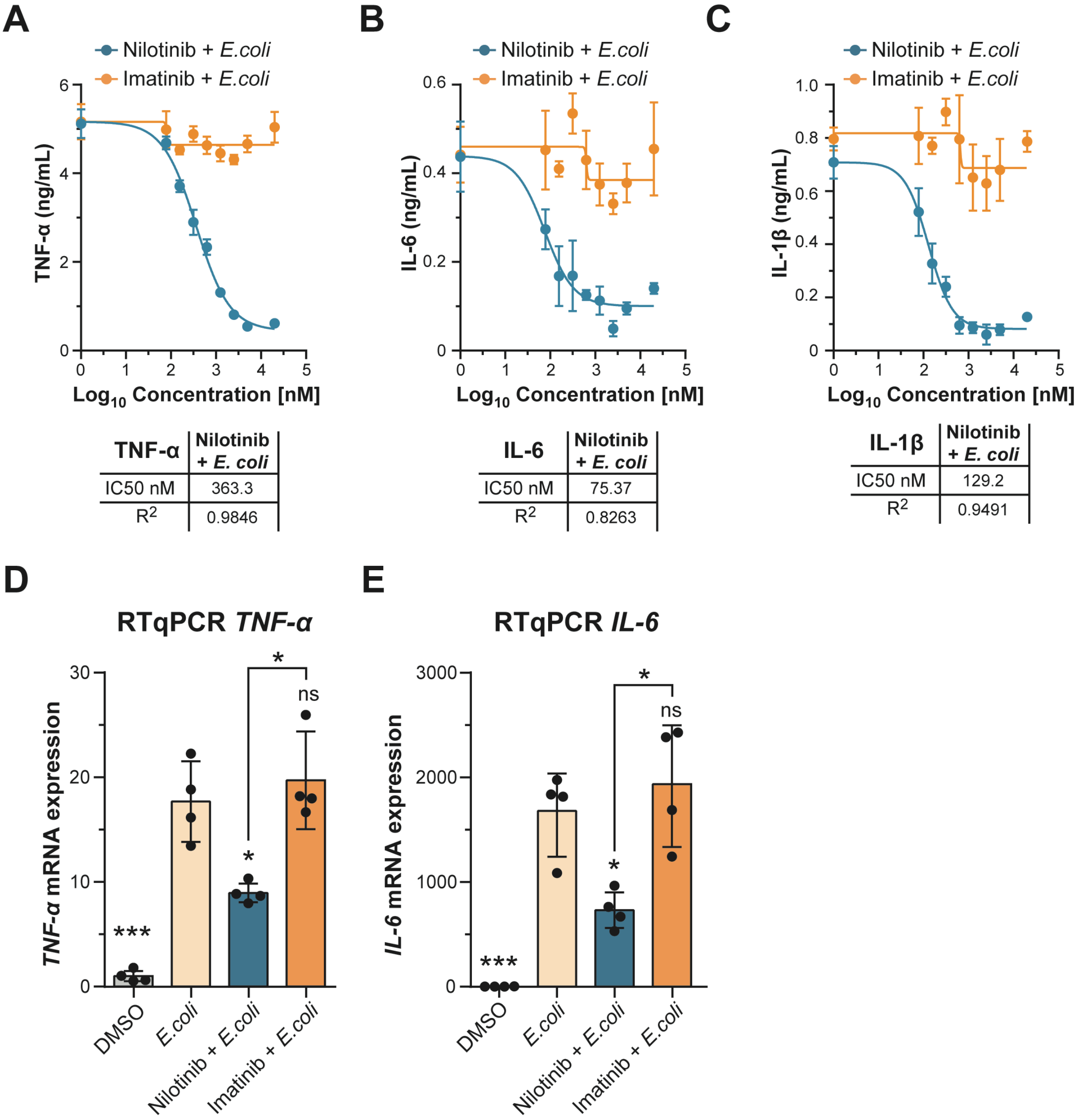


**Supplementary Figure 6. Nilotinib inhibits the pro-inflammatory phenotype in *E. coli*-activated monocytes. A-C)** ELISA IC50 curves of A) TNF-α, B) IL-6, and C) IL-1β were measured by ELISA at 24h. Error bars represent the standard deviation of four biological replicates. Significant differences between two groups were determined by a Students t-test. The statistical significance of the comparisons with *E. coli* is indicated as follows: ns, not significant; **, $P \leq 0.01$; ***, $P \leq 0.001$. Table insert shows IC50 (nM) and $R^2$. **D)** *TNF-α*, and **E)** *IL-6* expression were determined by RT-qPCR in THP-1 cells pre-treated with DMSO, 5 μM nilotinib or imatinib for one hour before stimulation with live *E. coli* for up to 24h. The results were analyzed using the $2^{-\Delta\Delta Ct}$ method and normalized using *GAPDH* and *TBP* as the reference genes; and DMSO as the reference sample. Error bars represent the standard deviation of four biological replicates. Significant differences between two groups were determined by a Mann-Whitney U-test. The statistical significance of the comparisons with *E. coli* is indicated as follows: ns, not significant; **, $P \leq 0.01$; ***, $P \leq 0.001$.

#### Supplementary Figure 7

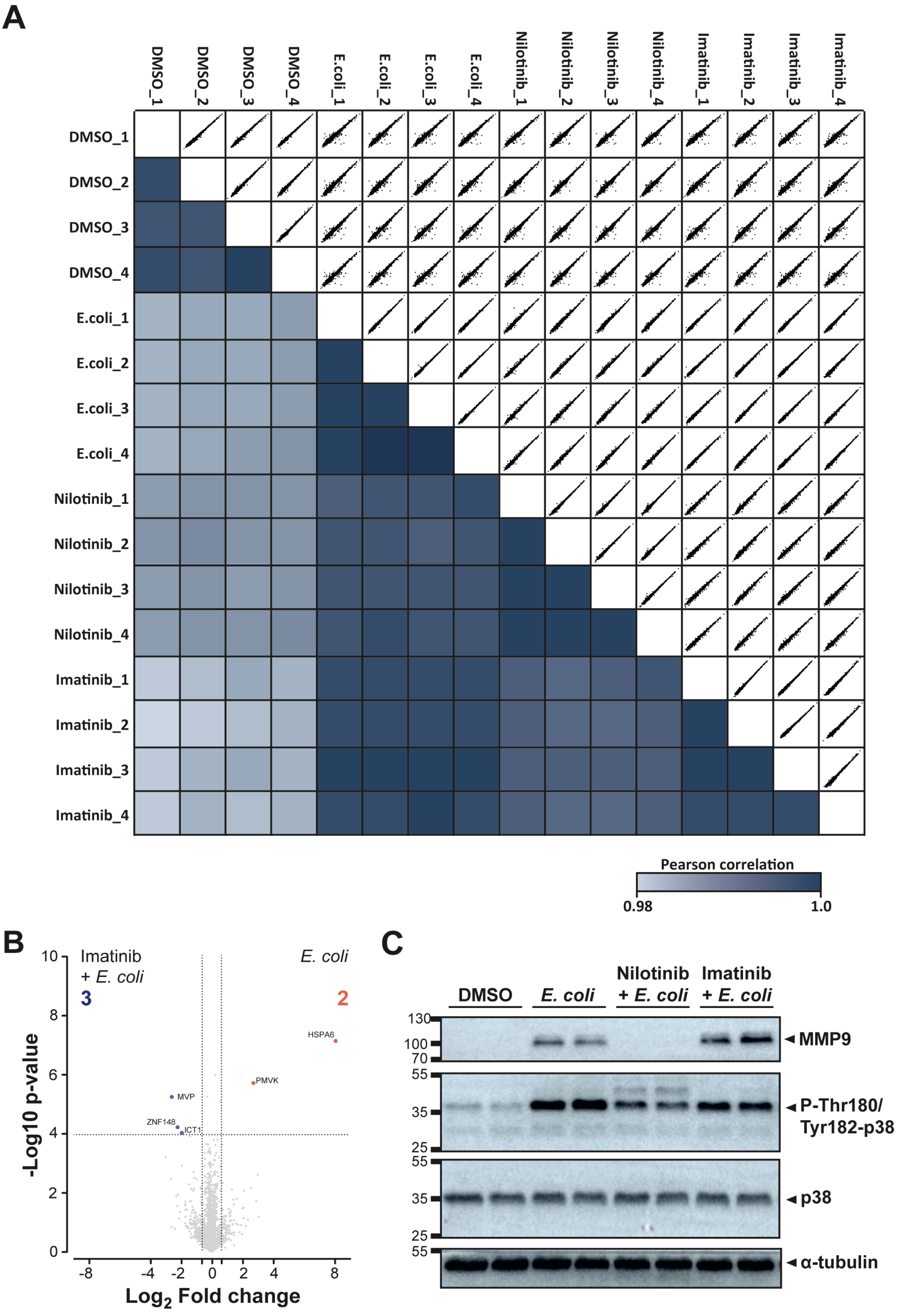

**Supplementary Figure 7. Differences in the proteome of nilotinib- and imatinib-treated monocytes. A)** The matrix of correlation plots revealed very high correlations between $\log_2$ transformed LFQ intensities (Pearson correlation coefficient between 0.982 and 0.998). The color code indicates the values of correlation coefficients. **B)** Volcano plot of THP-1 cells treated with 5 µM nilotinib *vs*. *E. coli*, cut-off of FDR <0.05 and 1.5-fold change between conditions. **C)** MMP9 and p38 levels in THP-1 pre-treated with 5 µM nilotinib or imatinib for one hour before stimulation with live *E. coli* for up to 24h. α-tubulin serves as a loading control. A representative image with two biological replicates of four replicates is shown. Relative mobilities of reference proteins (masses in kilo Daltons) are shown on the left of each blot.

#### Supplementary Figure 8

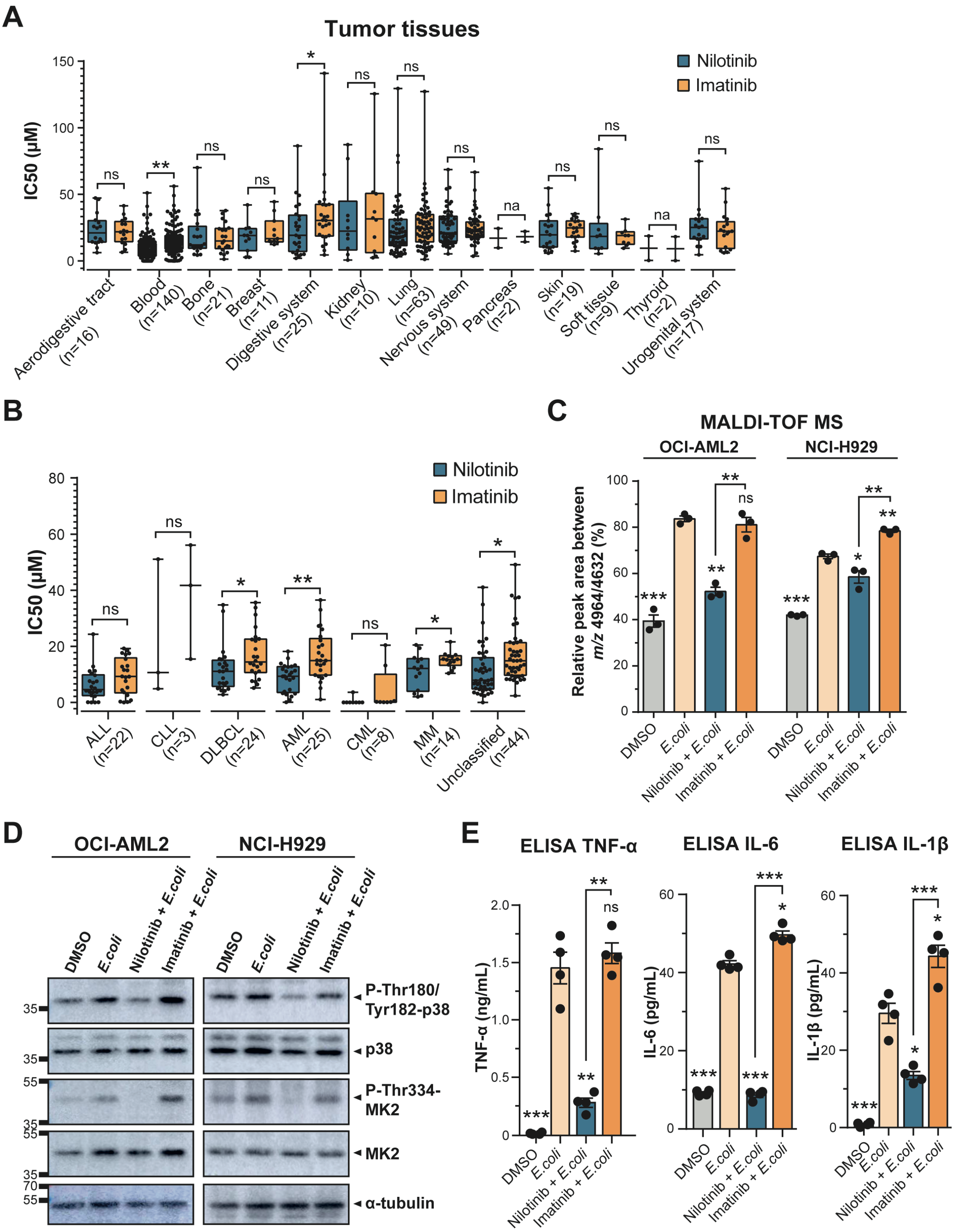


**Supplementary Figure 8. Nilotinib and imatinib show different sensitivity in blood malignancies. Nilotinib reduces the inflammation phenotype in OCI-AML2 and NCI-H929 cell line. A)** IC50 values of nilotinib and imatinib in various types of cancer, and **B)** in hematological malignancies according to TCGA classification from the Genomics of Drug Sensitivity in Cancer (GDSC) database. **C)** MALDI-TOF MS relative quantitation of the ratio m/z 4964/4632 in NCI-H929 cells pre-treated with DMSO, 5 µM nilotinib or imatinib for one hour before stimulation with live E. coli for up to 24h. **D)** p38 MAPK pathway status in NCI-H929 cells. α-tubulin was detected as a loading control. A representative image of four replicates is shown. Relative mobilities of reference proteins (masses in kilo Daltons) are shown on the left of each blot. **E)** TNF-α, IL-6, and IL-1β secretion was measured by ELISA at 24h. Error bars represent the SEM of four biological replicates. Significant differences between two groups were determined by a Mann-Whitney U-test. The statistical significance of the comparisons with *E. coli* is indicated as follows: ns, not significant; na, not applicable; *, $P \leq 0.05$; **, $P \leq 0.01$; ***, $P \leq 0.001$.
